# Precision neoantigen discovery using large-scale immunopeptidomes and composite modeling of MHC peptide presentation

**DOI:** 10.1101/2021.04.30.442203

**Authors:** Rachel Marty Pyke, Datta Mellacheruvu, Steven Dea, Charles Abbott, Simo V. Zhang, Nick A. Phillips, Jason Harris, Gabor Bartha, Sejal Desai, Rena McClory, John West, Michael P. Snyder, Richard Chen, Sean Michael Boyle

**Affiliations:** Personalis, Inc., Menlo Park, CA; Stanford University, Palo Alto, CA

## Abstract

Major histocompatibility complex (MHC)-bound peptides that originate from tumor-specific genetic alterations, known as neoantigens, are an important class of anti-cancer therapeutic targets. Accurately predicting peptide presentation by MHC complexes is a key aspect of discovering therapeutically relevant neoantigens. Technological improvements in mass-spectrometry-based immunopeptidomics and advanced modeling techniques have vastly improved MHC presentation prediction over the past two decades. However, improvement in the sensitivity and specificity of prediction algorithms is needed for clinical applications such as the development of personalized cancer vaccines, the discovery of biomarkers for response to checkpoint blockade and the quantification of autoimmune risk in gene therapies. Toward this end, we generated allele-specific immunopeptidomics data using 25 mono-allelic cell lines and created *Systematic HLA Epitope Ranking Pan Algorithm* (SHERPA™), a pan-allelic MHC-peptide algorithm for predicting MHC-peptide binding and presentation. In contrast to previously published large-scale mono-allelic data, we used an HLA-null K562 parental cell line and a stable transfection of HLA alleles to better emulate native presentation. Our dataset includes five previously unprofiled alleles that expand MHC binding pocket diversity in the training data and extend allelic coverage in underprofiled populations. To improve generalizability, SHERPA systematically integrates 128 mono-allelic and 384 multi-allelic samples with publicly available immunoproteomics data and binding assay data. Using this dataset, we developed two features that empirically estimate the propensities of genes and specific regions within gene bodies to engender immunopeptides to represent antigen processing. Using a composite model constructed with gradient boosting decision trees, multiallelic deconvolution and 2.15 million peptides encompassing 167 alleles, we achieved a 1.44 fold improvement of positive predictive value compared to existing tools when evaluated on independent mono-allelic datasets and a 1.15 fold improvement when evaluating on tumor samples. With a high degree of accuracy, SHERPA has the potential to enable precision neoantigen discovery for future clinical applications.

## Introduction

The concerted efforts of innate and adaptive immunity help maintain homeostasis and fight pathogen attacks. The innate immune system reacts quickly and is largely non-specific while the adaptive immune system is highly specific, typically takes a longer time to develop and is long lasting. Within the adaptive immune system, T cells survey the health of a cell by examining MHC protein complexes on the cell surface. All nucleated cells express MHC on their surface, and those cells presenting non-self and aberrant peptides are identified and eliminated. Although several factors are needed to mount an immunogenic response from CD8+ T cells, MHC Class I presentation of relevant peptides is a gatekeeping step (1–3).

Early *in vitro* and *in vivo* experiments evaluating binding characteristics of various peptide MHC pairs indicated that presented peptides had allele-specific motifs. *In vitro* experiments primarily attempted to measure binding affinity of specific peptides and their cognate MHC complexes using competitive binding assays in a hypothesis driven manner (4). Advances in liquid chromatography and mass spectrometry (LC-MS/MS) heralded a new era of large scale immunopeptidomics and the ability to learn the rules of binding and presentation in an unsupervised way (5). However, the assignment of peptides to their cognate alleles was problematic (6). Further advances in using genetically engineered cell lines expressing a single allele of interest, often at high copy number, helped circumvent this problem and produce unambiguous HLA peptidomes (7).

Algorithms to model MHC binding have progressed along with these technological advances. Early approaches relied exclusively on MHC-peptide binding affinity data (8). As immunopeptidomics data have become increasingly available, more groups have incorporated these data into prediction models. Several recent approaches employ multi-allelic immunopeptidomics data, including clustering-based deconvolution (9,10), iterative assignment (11), and direct modeling (12). Other efforts capitalize on the unambiguous nature of mono-allelic data (7,13,14). Furthermore, some algorithms aim to model MHC-peptide binding alone (10,15,16) whereas others extend to modeling antigen processing and surface presentation as well (7,12,13,17). Although there is disagreement on the best way to model MHC presentation, there is clear consensus that advances in mass spectrometry have enabled scalable data generation and significantly improved MHC-peptide binding and presentation prediction algorithms.

With the goal of creating an accurate MHC-peptide binding and prediction algorithm, we generated a high-quality dataset comprising 25 mono-allelic cell lines. All of the mono-allelic cell lines are distinct from previously published datasets due to their stable transfection and background cell line (K562). We also profiled five of the alleles that have not previously been profiled using mono-allelic immunopeptidomics technologies, expanding the known MHC binding pocket diversity and the allelic representation in underprofiled Asian, African and Middle Eastern populations; the lack of information for underrepresented groups is an area of high current interest (18). Further, we combined our dataset with curated publicly available binding affinity data and mono-allelic immunopeptidomics data to create a robust MHC binding prediction algorithm. In order to capture the diverse facets of antigen processing and presentation, we systematically reprocessed and deconvoluted publicly available multi-allelic data from several tissue types with different transcript expression profiles using our binding model trained on mono-allelic data and expanded our training data to encompass 167 alleles. We also employed this large dataset to empirically learn patterns in antigen processing at the protein and peptide level. We applied gradient boosted decision trees to train our prediction models, which we call SHERPA (*Systematic HLA Epitope Ranking Pan Algorithm*). We evaluated SHERPA on held-out mono-allelic and independently generated multi-allelic tumor datasets to demonstrate that our integrated approach for MHC-peptide presentation prediction has a 1.15 fold improvement in performance over the best existing tools.

## Experimental procedures

### Immunopeptidomics using mono-allelic cell lines

#### Experimental design and statistical rationale

In this study, all peptides studied were derived from the MHC-I immunopeptidome. Experimental work was performed on two sample types: mono-allelic cell lines and tumor tissue. For the mono-allelic cell lines, 28 cell lines of 5×10^9^ cells were processed. Three biological replicates were assessed for a single cell line. One cell line was an HLA-null line used as a negative control. Peptides from 8 of the cell lines were processed a single time. Peptides from the other 20 of the cell lines were divided and processed on the mass spectrometer twice (11 with CID only, 9 with both CID and EThcD). For the tumor tissues, 12 tissues were processed without controls or replicates. In addition to the experimental work, publicly available peptides were also analyzed. A Wald test and two-sided T tests were used in this study.

#### Cell culture

We generated mono-allelic cell lines by stably transfecting K562 parental cells with a single allele of interest using Jump-In™ technology (Thermo Fisher Scientific). Optimized plasmids comprising the sequences for HLA, beta 2 microglobulin (B2M) and IRES promoter were synthesized for each allele (GeneArt). Cells were screened for plasmid integration and expanded to 500M cells through various passages. The cells were pelleted once surface expression of target alleles was confirmed using flow cytometry (using W6/32 antibody). Transfection experiments were performed by Thermo Fisher Scientific.

#### Immunoprecipitation of peptide-MHC complexes

Pelleted cells were resuspended in octylthioglucoside lysis buffer and the cell lysate was incubated overnight with W6/32 antibody immobilized on Protein A sepharose. After washing the resin, MHC bound peptides were eluted using 0.1 M acetic acid, 0.1% TFA. The success of immunoprecipitation was verified by confirming the depletion of MHC Class I complex in post-IP samples using ELISA (enzyme-linked immunosorbent assay). Immunoprecipitations were performed by Cayman Chemical.

#### Peptide sequencing using LC-MS/MS

Eluted peptides were desalted using solid phase extraction (SPE; Empore C18), loaded on a column, eluted using 80/20 acetonitrile/water (0.1% TFA), lyophilized and stored. Samples were reconstituted in 0.1% TFA before they were analyzed using liquid chromatography mass spectrometry (LC/MS/MS). Chromatographic separation was performed using a 2h gradient on a Waters NanoAcquity system. Peptides were analyzed using a ThermoFisher Fusion Lumos mass spectrometer in data dependent mode (MS1: Orbitrap at 60,000 FWHM resolution, m/z range: 300-800; isolation window: 1.6 Da; fragmentation: EThcD and CID; MS2: Orbitrap at 15,000 FWHM; cycle time: 3s). Mass spectrometry experiments were performed by MS Bioworks, LLC.

#### Peptide identification

Peptides were identified using PEAKS software (PEAKS Studio 10.0 build 20190129) (19) using the default two step identification workflow, where the first step performs de novo sequencing to identify mass tags and the second step performs a database search on a subset of putative proteins identified using de novo mass tags. The workflow was run with the following settings. Protein database: Swissprot proteome database (20,402 entries; dated 03-20-2019); precursor mass tolerance: 10 ppm; Fragment mass tolerance: 0.02 Da; Enzyme specificity: none; Fixed modifications: carbamidomethylation of cysteine (+57.0215); Variable modifications: oxidation of Methionine (+15.9949), N-terminal acetylation (+42.0106). Peptide-to-spectrum matches (PSMs) were filtered at 1% FDR, estimated using decoy sequences.

#### Post processing and quality control

Peptides identified at 1% false discovery rate (FDR) were filtered further to remove spurious peptides using the following constraints: spurious peptides were defined as polymeric peptides, peptides from highly chimeric spectra (n>2) and identifications with less than 10 fragment ions (**Supplementary Code**). Background contaminants, profiled using a mock transfection (GFP), were filtered out. Samples were manually inspected for the presence of motif signatures (using Gibbs Clustering software (9)) and peptide yields (heuristic cutoff of 500 peptides per sample).

### Extraction and processing of public data sets

#### Extraction and processing of publicly available mono- and multi- allelic data

Publicly available immunopeptidomics data were identified after an exhaustive literature search. Raw data (.raw files) were downloaded and systematically processed similar to in-house data. HLA types to sample mappings were extracted from corresponding publications. Peptide identifications from samples that passed rigorous quality control criteria (as described above) were aggregated (**Supp. Table 2**).

#### Processing of in vitro binding affinity data

Raw HLA-peptidome data were downloaded from the Immune Epitope Database (IEDB) (date: 03-09-2020) and filtered to identify entries corresponding to *in vitro* binding assays. The ‘Object Type’ column was filtered to exclude any non-linear peptides and the ‘Units' column was filtered to exclude any non-nM entries. Further, only four digit MHC class I peptides of length 8-11 were retained. The peptides were derived from the following ‘Method/Technique’ categories: ‘purified MHC/competitive/radioactivity’, ‘purifiedMHC/direct/fluorescence’, ‘purified MHC/competitive/fluorescence’, ‘cellularMHC/competitive/fluorescence’,cellular MHC/direct/fluorescence’, ‘cellular MHC/competitive/radioactivity’, ‘binding assay’ and ‘lysate MHC/direct/radioactivity’. Ligands with IC50 values less than 500 nm were identified as binders (**Supp. Table 2**).

### Measurement and analysis of transcript expression

#### Sequencing of mono-allelic cell lines

Representative samples of mono-allelic K562 cell lines generated in-house were sequenced in triplicate using our in-house commercial platform ImmunoID NeXT™ with 200 million paired end reads (150 base pair) of sequencing for RNA from tumor samples. Reads were aligned in accordance with Personalis Cancer RNA pipeline and transcript per million (TPM) values were extracted. Another cell line, B721.221, with large amounts of publicly available mono-allelic data, was sourced from the American Type Culture Collection (ATCC) and profiled in triplicate using ImmunoID NeXT similarly to minimize technological variance.

#### Generation of transcriptome for external immunopepdomics datasets

For one of the largest publicly available mono-allelic datasets generated using B721.221 cell (MSV000080527; MSV000084172) lines, we re-generated the transcriptome data on ImmunoID NeXT (7,13), as described above. For the rest of the samples in the expanded dataset (**Supp. Table 2**), we imputed TPM values with our internal database by taking the median values from all samples with a matching tissue type.

#### Evaluation of differential transcriptome abundances across cell lines and tissue types

To evaluate the differences in expression between K562 and B721.221, we performed differential gene expression analysis using the Deseq2 package (20), which takes in raw transcript counts from all replicates of a sample set to determine differentially expressed genes. We used a threshold of an absolute value log_2_ fold change of greater than or equal to 2 in conjunction with an adjusted p-value threshold of less than or equal to 0.01 to identify differential gene expression. We then visualized the differentially expressed genes in a volcano plot to show cell line gene expression differences, using the Bioinfokit tool by Renesh Bedre (https://github.com/reneshbedre/bioinfokit/tree/v0.9). We then took the identified genes that were enriched in K562 and the genes that were depleted in K562 as compared to B721.221 and calculated the Gene Ontology (GO) enrichment terms for both gene sets using http://geneontology.org/ (**Supp. Table 3**). To evaluate the differences in expression between the tissue types, we restricted our analysis to the tissues with unique expression profiles (n=30). Then, we identified the 5,000 genes with the highest standard deviation between the expression profiles and visualized those genes as rows and the unique expression profiles as columns in a clustered heatmap.

### Exploratory analysis of immunopeptidomics data

#### Relationship between expression and peptide presentation

Peptides identified in K562 mono-allelic samples were assigned a transcript expression value (TPM) by mapping each peptide to one or more proteins, and proteins to one or more transcripts. The highest TPM was assigned as the representative value when there are multiple mappings. A set of random peptides (n >50 million) were also generated and assigned TPMs as described. TPMs were binned to deciles and enrichment of presented peptides in each bin was calculated as the log ratio of number of presented peptides to background random peptides.

#### Relationship between cleavage specificities and peptide presentation

To understand the cleavage preferences of the proteasome in our mono-allelic cell lines, we mapped each peptide to a protein (as described above) and identified the right and left flanking sequences (five amino acids) of the peptides. A set of random peptides (n >50 million) were also generated and assigned flanking sequences as described to serve as a null dataset. To calculate the enrichment or depletion of amino acids at each position in the flanking regions, we subtracted the observed amino acid frequency from the random amino acid frequency and divided by the random amino acid frequency. Of note, the c- and n-terminus of the protein was also considered in addition to the individual amino acids. We visualized the enrichment and depletion as a heatmap.

#### Clustering alleles by binding pocket similarity

The binding pocket was represented by a pseudo sequence of 34 amino acids as described previously (8). The distance between alleles was calculated by taking the sum of the BLOSUM62 scores for the amino acids in each allele at each position across the pseudo sequence. All of the unique alleles with in-house or public mono-allelic immunopeptidomics data were compared with one another and visualized as a heatmap.

#### Estimating allelic coverage in underrepresented ethnic populations

Allele frequencies from several ethnic populations were downloaded from http://www.allelefrequencies.net/ on 05-22-2020. The frequencies of the five novel alleles in underrepresented ethnic populations were taken directly from the data on the website. To estimate the allelic coverage of all of the unique alleles in the whole data across ethnic populations, we focused on the data derived from the National Marrow Donor Program (NMDP, n=18 populations). For each population, we performed a monte carlo simulation to generate 10,000 synthetic individuals with the allelic frequencies for that population specified by NMDP. Then, we calculated the percentage of the alleles in the synthetic cohort that were represented in the expanded dataset (in-house mono-allelic, public mono-allelic, IEDB and public multi-allelic).

#### Generation of peptide motifs

We generated the motifs for each allele using the weblogo software (version: 3.7.1). All motifs were based on a specific length of peptide (8-, 9-, 10- or 11-mers). Motifs for public datasets were based on the peptides derived from our internal processing pipeline.

#### Generation of position-specific frequencies of amino acids of the binding pocket

To assess the representativeness of our expanded dataset to the full space of known alleles, we calculated the frequency of amino acid at each position of the binding pocket pseudo sequence for both all of the alleles in the IMGT (the international ImMunoGeneTics information system; http://www.imgt.org/) and our expanded dataset. We visualized these frequencies as stacked bar plots.

### Feature engineering and building prediction models

#### Data splitting for training and evaluation

In order to rigorously evaluate our performance on novel peptides, we ensured that no peptides with overlapping 8-mer cores were present across the training, validation and testing datasets. We took the unique set of peptides across our mono-allelic immunopeptidomics, multi-allelic immunopeptidomics and IEDB datasets. Then, we grouped peptides that contained any identical 8-mer substrings (“nested” peptides) and placed each peptide group in one of ten different subsets. Once we had roughly equal peptide numbers in each subset, we assigned one subset to be for validation (~10%), one subset to be for testing (~10%) and the final eight subsets to be for training (~80%). Moreover, we fully held out 32 multi-allelic samples from the training dataset (described in detail in the ‘Benchmarking and evaluation of prediction models' section) and evaluated the models on a subset of the peptides (~10%) that were fully held out from training (as described above). All publicly available multi-allelic data (except for the 32 held out samples) was used for feature engineering of the gene propensity and hotspot scores because systematically holding out proteins or peptides would skew the scores. Only the ~80% training subset of the multi-allelic data was used for deconvolution.

#### Specifications of prediction models

We trained two types of prediction algorithms, one that models *binding* and another that models *presentation*. The binding algorithm models MHC peptide interaction affinities and takes as inputs the HLA binding pocket, the amino acid sequence and the length of peptide ligands. The presentation algorithm jointly models both antigen processing and peptide-MHC (pMHC) binding, whose inputs include: the HLA binding pocket, amino acid sequence and length of peptide ligands, proteasomal processing footprints manifested in the left and right flanking regions of peptide ligands, abundance of source proteins engendering peptide ligands as measured by gene expression, and two features that model the propensity of antigen processing and presentation. Features corresponding to these inputs as generated as follows:

1. *HLA binding pocket (B):* The binding pocket is represented by a pseudo sequence of amino acids as described previously (8). Briefly, 34 positions on the protein sequence of the HLA that are a distance of 4 Å or lesser in crystallographic structures were selected from the full protein to serve as the pseudo sequence. The amino acid sequence is encoded using a BLOSUM62 substitution matrix, where each amino acid is represented by a 20-dimensional vector constituting the relative weights of amino acid substitutions. We chose BLOSUM62 encoding as opposed to a one-hot encoding approach under the assumption that substitutions between amino acids with evolutionary similarity have a lower impact on epitope binding changes than substitutions between amino acids that are very dissimilar.
2. *Peptide ligand (P):* The amino acid sequence of the peptide ligand is encoded using a BLOSUM62 substitution matrix, where each amino acid is represented by a 20- dimensional vector constituting the relative weights of amino acid substitutions. Since HLA ligands could be of variable lengths (8- to 11-mers), we adapted a middle-padding approach described previously and adjusted all peptides to a length of 11 amino acids by inserting blanks in the middle (12). Briefly, for every peptide with fewer than 11 amino acids, we assign one to three 20-dimensional vector(s) of zeros to the center of the peptide encoding to reach the maximal 11-mer length. We take this approach to have a pan-length algorithm that keeps the peptide anchors in consistent columns of the matrix.
3. *Peptide length (L):* The number of amino acids in the peptide ligand is designated as the peptide length.
4. *Left and right flanking regions (F):* Peptide sequences (5-mers) to the left and right of the peptide ligand in the source protein are used as the left and right flanking regions respectively. Multi-mappers are resolved by assigning the protein with the highest transcript expression to the peptide. These left and right flanking 5-mers are encoded using BLOSUM62 substitution matrix as described above.
5. *Abundance of source protein (T):* Peptide ligands are redundantly assigned to all source proteins and then to transcripts. The transcript with the highest expression (calculated as the TPM) is chosen. Both the transcript and the TPM are then assigned to the peptide ligand.
6. *Gene propensity score (G): P*ublicly available multi-allelic data was used to estimate gene propensity. Peptide ligands from each sample are mapped to source proteins and then to transcripts redundantly. The number of peptides mapping to each transcript-associated protein were determined. To calculate the expected number of peptides for each transcript-associated protein, the number of transcripts per million for each protein across all multi-allelic data sources are determined and normalized by the protein length and the number of peptides derived from each sample. Then, the number of observed and expected peptides for each protein are summed across all of the multi-allelic datasets. Finally, the observed values were divided by the expected values to derive a gene propensity score for each protein (gene). Proteins without any observed transcripts across all samples are (potentially pseudo genes) given a score of −3 to deprioritize them.
7. *Hotspot score (H):* Publicly available multi-allelic data were used to estimate the hotspot score. Peptide ligands from each sample are mapped to source proteins. When peptides mapped to multiple proteins, all potential mappings are used. The number of peptides overlapping a particular amino acid represents the hotspot score of that amino acid. To assign a hotspot score for a peptide, the peptide is mapped to its source protein and the amino acid hotspot score spanning the peptide is averaged.

We evaluated the contribution of each feature to the XGBoost model using the ‘gain’ metric, which measures the relative contribution of a feature to the main model by aggregating the individual contributions of the feature in each tree. Higher values indicate greater importance.

#### Generating negative examples (non-binders)

Immunopeptidomics experiments only generate peptides that successfully bind to and are presented by MHC molecules. Thus, we synthetically generated negative examples instead of experimentally identifying them. We generated 20 negative examples for every positive example in our training and validation datasets. To generate a negative example, we randomly selected a protein from the Swissprot proteome (downloaded on 03-20-2019) and then randomly selected a peptide from within that protein. Peptides were selected to have length 8, 9, 10 or 11 with equal probability. Flanking regions were assigned based on the true flanking regions around the selected peptides. A gene expression value (TPM) was assigned by randomly selecting a transcript from the transcriptome of the associated positive example. The gene propensity score and hotspot score were assigned based on the protein and position in the protein of the peptide selected for the negative example.

#### Training the prediction algorithm

All models were trained using a gradient boosted decision tree algorithm implemented using an open-source package XGBoost (21). All numeric and encoded features were provided as a vectorized input feature vector for training the algorithm. Optimal training parameters were selected based on a subset of samples using sequential model-based optimization, implemented using the open-source package HyperOpt (22). The resulting training parameters used for the final training were as follows: loss function - binary logistic; max depth - 10; eta - 0.01; subsample - 0.7; early stopping rounds - 5; min child weight - 0.5; max delta step - 1; tree method - hist; number of estimators - 500. Large training sets were subset and processed in parallel based on available compute resources and processed on a high-performance cluster.

#### Calibrating raw scores using percent rank values

A set of 500,000 peptides were randomly selected from the human proteome. Once we trained a model, we calculated the predictions across the set of random peptides with every allele. Then, we ranked the random peptides according to their raw prediction probability (output of XGBoost) for each allele. For each new peptide predicted, the assigned rank is the percentage of the random set that is predicted to bind or to be presented with a better raw score than the new peptide. The ranks range from 0 to 100 with lower scores meaning better bound or presented peptides. The ranks are re-calculated for each model and allele combination.

#### Applying prediction models to multi-allelic cell line and patient samples

The MONO-Binding model was used to generate binding ranks for the peptides in the public multiallelic samples (including tumors, tissues and cell lines) with immunopeptidomics data. Only peptides with 8, 9, 10 and 11 amino acids were considered. 10% of multi-allelic data was held out for testing purposes. Predictions were made for all HLA alleles (up to six) of each sample. Samples without HLA typing were excluded from this analysis.

#### Deconvolution of multi-allelic data

Once predictions were made for all HLA alleles of each multi-allelic sample with immunopeptidomics data, the data was deconvoluted to decide which allele-peptide pairs should be included in the training dataset for the final model. First, we excluded all allele-peptide pairs with a predicted binding rank of ≥ 0.5 to exclude all peptides that do not bind to any of the designated alleles. Second, if there were multiple alleles predicted to bind to a specific peptide, we selected the allele-peptide pair with the lowest rank (best binder) and excluded all other pairs. Then, we removed any duplicate allele-peptide pairs. Finally, we generated 20 negative examples (as described above) for every new positive example derived from the multi-allelic data.

#### Training composite models

We trained a total of three prediction models for our composite model. A schematic of the composite model is shown in **Figure 5A** and the details are outlined in **Supp. Table 4**. In addition, we also trained five more models to help us better understand features that contribute to optimal performance. The prediction models in our composite model were trained as follows:

1. *MONO-Binding*: Trained using the IEDB, in-house mono-allelic immunopeptidomics and public mono-allelic immunopeptidomics data with B, P and L as features.
2. *SHERPA-Binding*: Trained using the IEDB, in-house mono-allelic immunopeptidomics, public mono-allelic immunopeptidomics and deconvoluted multi-allelic immunopeptidomics data with B, P and L as features.
3. *SHERPA-Presentation*: Trained using the in-house and public mono-allelic immunopeptidomics data with SHERPA-Binding, F, T, G and Has features.

The additional prediction models were trained as follows:

1. *PUBLIC-Binding*: Trained using the public mono-allelic immunopeptidomics data with B, P, and L as features.
2. *MONO-Binding-LOO*: 126 allele-specific models trained using the IEDB, in-house mono-allelic immunopeptidomics and public mono-allelic immunopeptidomics data with B, P, L, IEDB-Binding, INHOUSE-Binding and PUBLIC-Binding as features. Each allele-specific model was trained without peptides in the training dataset from the respective alleles.
3. *SHERPA-Binding*+*F*: Trained using the IEDB, in-house mono-allelic immunopeptidomics, public mono-allelic immunopeptidomics and deconvoluted multiallelic immunopeptidomics data with SHERPA-Binding and F as features.
4. *SHERPA-Binding*+*FT*: Trained using the IEDB, in-house mono-allelic immunopeptidomics, public mono-allelic immunopeptidomics and deconvoluted multiallelic immunopeptidomics data with SHERPA-Binding, F and T as features.
5. *SHERPA-Binding*+*FTG*: Trained using the IEDB, in-house mono-allelic immunopeptidomics, public mono-allelic immunopeptidomics and deconvoluted multiallelic immunopeptidomics data with SHERPA-Binding, F, T and G as features.

### Benchmarking and evaluation of prediction models

#### Generation of the mono-allelic held-out test data

For the mono-allelic immunopeptidomics datasets (in-house and public), ~10% of the positive examples were held out of the training and validation datasets. Each positive example was supplemented with 999 negative examples to reflect the positive to negative ratio accepted by the field (5,23–26). For the IEDB data, ~10% of the positive and negative data was withheld from the training dataset. No supplementary negative examples were added, so the ratios in the test dataset reflect the positive to negative peptide ratio in IEDB. All mono-allelic validation figures rely on the mono-allelic immunopeptidomics test data except for **Supp. Figure 8B**, which uses the IEDB test data.

#### Calculation of evaluation metrics

Three metrics were used to evaluate the performance of the prediction models. The metrics are as follows:

1. *Positive predictive value (PPV)*: PPVs were calculated by making predictions on the entire test dataset and calculating the percentage of peptides in the top X% of predictions that are positive examples, with X representing the portion of the dataset that are positive examples. Each PPV was calculated individually for the peptides of each allele and combined using a median giving a single metric. Of note, the positive to negative ratios of the mono-allelic immunopeptidomics and IEDB test datasets vary, so the interpretations of the plots are different.
2. *Precision-recall curves*: Precision-recall curves were generated by calculating the precision and the recall for every possible cutoff and plotting them as a single line.
3. *Fraction of observed peptides predicted by model*: This metric was used for all multi-allelic tumor validation analyses. First, a single score is selected to represent each peptide observed with immunopeptidomics by making predictions on all of the patient's HLA alleles and selecting the best (lowest) rank among the predictions. Then, the score is calculated by determining the percentage of observed peptides that are given a rank of ≤ 0.1.

#### Leave-one-out pan-allelic analysis

To evaluate the pan-allelic performance of the MONO-Binding model used for model-based deconvolution, we trained 126 independent models with the same features as the MONO-Binding model. For each model, we excluded the peptides from a specific allele from the set of peptides used to train the MONO-Binding model. To generate the predicted motif for each allele **(Supp. Figure 6)**, we predicted the binding rank for 500,000 random peptides for a given allele with the model for which the allele had been excluded from training. Then, we generated the motif for peptides with the top percentile of binding ranks. Motifs were only visualized for alleles with at least 50 positive peptides in the training data. To generate the precision recall curve, we used all 126 models to predict the binding ranks of the mono-allelic immunopeptidomics data (in-house and public) that was excluded from training and validation (10% test dataset). The predictions for each allele were made with the model that excluded that allele from training.

#### Generating validation data using tissue samples

A total of 12 fresh frozen tumor samples (five colorectal and seven lung) and matched adjacent normal fresh frozen samples were selected for patient validation. These samples were purchased from a biobank and were collected under IRB-approved protocols and abide by the Declaration of Helsinki principles. Each tumor sample was divided into two pieces. Immunoprecipitation of MHC complexes followed by LC-MS-MS (as described above) was performed in a portion of each tumor sample to yield immunopeptidomics data. DNA and RNA was extracted from the remaining tumor sample and DNA was extracted from the adjacent normal sample for analysis with ImmunoID NeXT (as described above). The RNA extracted from the tumor sample was used to yield transcriptomic data.

#### Extraction of held-out validation data sets from external multi-allelic samples

~10% of multi-allelic immunopeptidomics data with non-overlapping ‘nested’ peptides with the training and validation datasets was withheld from deconvolution and training to serve as a test dataset. We used 10 samples from each of two specific datasets to validate our internal tumor immunopeptidomics performance (PXD007635, PXD009602) (27,28). Only samples with HLA-A, -B and -C typing at 4-digit resolution were used for the analysis.

#### Immunogenicity evaluation

In order for a peptide to incite an immunogenic response, it must be presented on the cellular surface by an MHC allele. Thus, we evaluated the ability of the algorithms to positively identify immunogenic peptides. We used the dataset described in Chowell et al for this analysis and focused exclusively on the immunogenic peptides (29). Then, we evaluated the percentage of immunogenic peptides that were predicted by the various models at <= 0.1 percentile rank.

#### Running comparison prediction algorithms

NetMHCpan-4.1-BA, NetMHCpan-4.1-EL and MHCFlurry-2.0-BA were used as comparison prediction algorithms. All algorithms were run according to default settings. Percentile rank outputs were used for each analysis.

## Results

### Generation of mono-allelic immunopeptidomics data

In order to generate high-quality data, we used an HLA-null K562 parental cell line and engineered mono-allelic cell lines that expressed a single HLA allele of interest (**Figure 1A**). Expanded cells with good surface expression of target alleles were used for immuno-precipitation of peptides associated with MHC complexes, followed by elution of ligands and peptide sequencing using liquid chromatography mass spectrometry (LC-MS/MS). Our stable transfection protocol, which ensures single-site integration of the target allele, minimizes expression biases and enables modeling of antigen processing in closer-to-native condition. Moreover, our K562 parental cell line provides a different background than the majority of previously published mono-allelic data (B721.221 parental cell line), providing a novel and standardized system to study peptide presentation.

**Figure 1:**
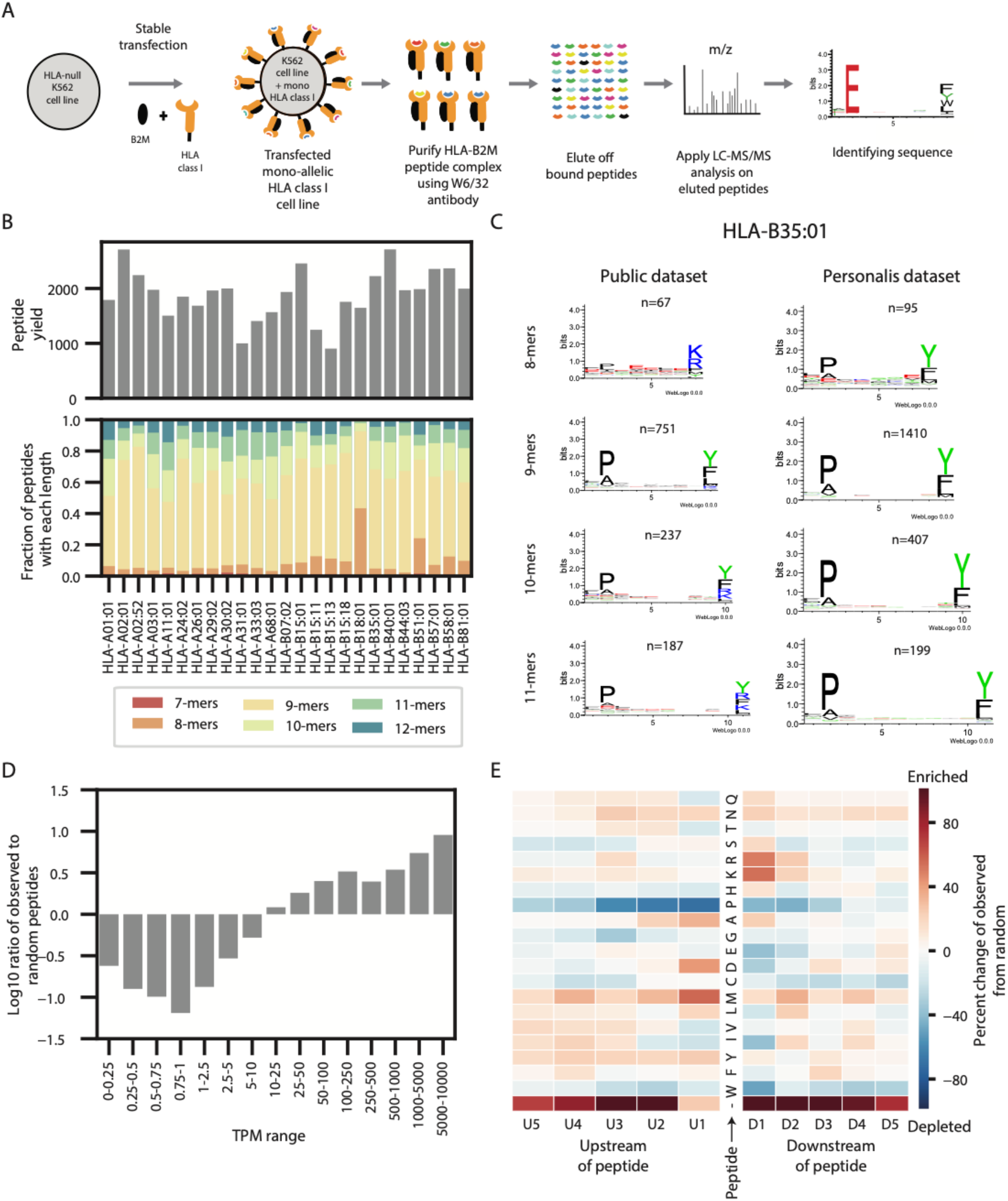
Generation and overview of the mono-allelic data. **(A)** A schematic of the experimental procedure to generate the mono-allelic training data. An HLA allele and B2M were stably transfected into an HLA-null K562 parental cell line. The MHC-peptide complex was purified using a w6/32 antibody and the peptides were gently eluted off the complexes. The peptides were sequenced with LC-MS/MS and identified with a database search. **(B)** Bar plots showing the peptide yields and distribution of peptide lengths for each of the 25 mono-allelic cell lines. **(C)** A comparison between motifs of peptides generated from our mono-allelic cell line with HLA-B35:01 and a publicly available dataset for the same allele. Motifs are shown for peptides of length 8, 9, 10 and 11. See Supplementary Figure 1 for motifs from all 25 cells. See Supplementary Figure 2 for comparisons with other public datasets. **(D)** A bar plot showing the distribution of the ratio of observed peptides from the mono-allelic cell lines compared to random expectation across several TPM ranges. Values are shown with a log10 transformation. **(E)** A heatmap showing the enrichment and depletion of five amino acids upstream and downstream of the peptides identified from the mono-allelic cell lines compared to a random expectation. Red denotes the enrichment of amino acids and blue denotes the depletion of them. The c- and n- terminus of the protein are denoted with ‘-’.

We generated data for 25 alleles (**Supp. Table 1**) and identified an average of 1,575 unique peptides per allele (range: 905-2712) (**Figure 1B**). We first evaluated the allele-wise length distribution and recapitulated previously described observations, with HLA-A alleles presenting longer peptides than HLA-B alleles (Two-sided t-test, p=0.0066, **Supp. Figure 1A**) (13). We also compared the motifs in our raw data to those published previously. Whereas most of the motifs are concordant, we also observed some differences (**Supp. Figure 2**). As an example, a previously published dataset (7,13) for HLA-B*35:01 shows a tryptic digest like lysine/arginine signature for 8, 10 and 11-mers on the c-terminal end of the peptide, whereas our data shows a motif that is consistent with the ones observed with 9-mers (**Figure 1C**). We observe similar patterns in HLA-A*29:02, HLA-A*30:02, and HLA-B*57:01 (**Supp. Figure 3**), suggesting that there may be contamination in the publicly available datasets. Since MHC binding algorithms learn allele-specific motifs, such subtle differences are critical to create accurate prediction models. We then evaluated the relationship between transcript abundance and peptide presentation (**Figure 1D**). We observed a positive correlation between the propensity of a transcript to engender presented peptides and its expression level, represented by the transcript per million (TPM). We also evaluated the proteasomal cleavage signatures of presented peptides by examining the left- and right-flanking amino acids (**Figure 1E**). The relative enrichment of amino acids compared to a random background set indicates an overrepresentation of lysine and arginine on the C-terminal end of the peptide. This observation is in concordance with previously described tryptic and chemo-tryptic enzymatic activity of proteasome cleavage (7,13). Of note, we observe an enrichment of peptides originating from either the C-terminal or N-terminal of the source protein, indicating preferential proteasomal processing of peptides requiring relatively lesser enzymatic cleavage (Two-sided t-test, p=3.77e-32, **Supp. Figure 1B**). Enrichment was not observed in the N-terminal end of the peptide in other mono-allelic datasets (7,13), possibly due to overexpression of MHC with transient transfection protocols leading to a lack of discrimination among presented peptides.

### Novel mono-allelic cell lines enhance diversity of training data and reveal subtle binding preferences

Beyond the added value of our unique parental cell line and a stable transfection approach, our dataset includes five alleles that had not previously been profiled using mono-allelic cell lines. We selected these five alleles on the basis of two criteria: (1) the uniqueness of the allele binding pocket compared to previously profiled alleles (**Figure 2A**) and (2) the population frequency of the allele in ethnic populations that are underrepresented in current datasets (**Figure 2B**). For example, we clustered alleles by their binding pocket similarity and selected HLA-A*02:52 from the HLA-A*02 subcluster due to its high frequency in Iranian Kurdish populations (7% frequency, **Figure 2B**). Though several HLA-A*02 alleles have been profiled by previous groups (light grey), we found small differences in the motif from closely clustered alleles despite very similar binding pockets. For example, unlike surrounding HLA*02 alleles, we observed the presence of tyrosine in the second anchor position (C-terminal end of peptide) for HLA-A*02:52. Moreover, although HLA-A*02:52 clusters tightly with HLA-A*02:07, HLA-A*02:07 has a strong preference for aspartic acid at position three whereas HLA-A*02:52 has no preference. Similarly, we profiled three HLA-B*15 alleles (HLA-B*15:11, HLA-B*15:13 and HLA-B*15:18) that are frequent across several Asian populations, including the second most frequent allele in a Beijing population at over 12% (HLA-B*15:11). Despite inclusion in the same allele sub-group and having binding pockets that cluster together, the three alleles have very distinct associated peptide motifs with each having different preferences at each anchor position. These three alleles highlight the value of our novel alleles in the creation of a pan-allelic model which relies on learning how amino acid changes impact binding preference. Another novel allele, HLA-B*81:01, falls outside of a major binding pocket cluster, but its peptide motif shows strong similarities to surrounding alleles (**Figure 2A**). Of note, HLA-B*81:01 has moderate population frequency across over a dozen African populations (2-6% frequencies). Taken together, our novel alleles enhance both binding pocket diversity and population representation in a manner that is not represented in previously published training data.

**Figure 2:**
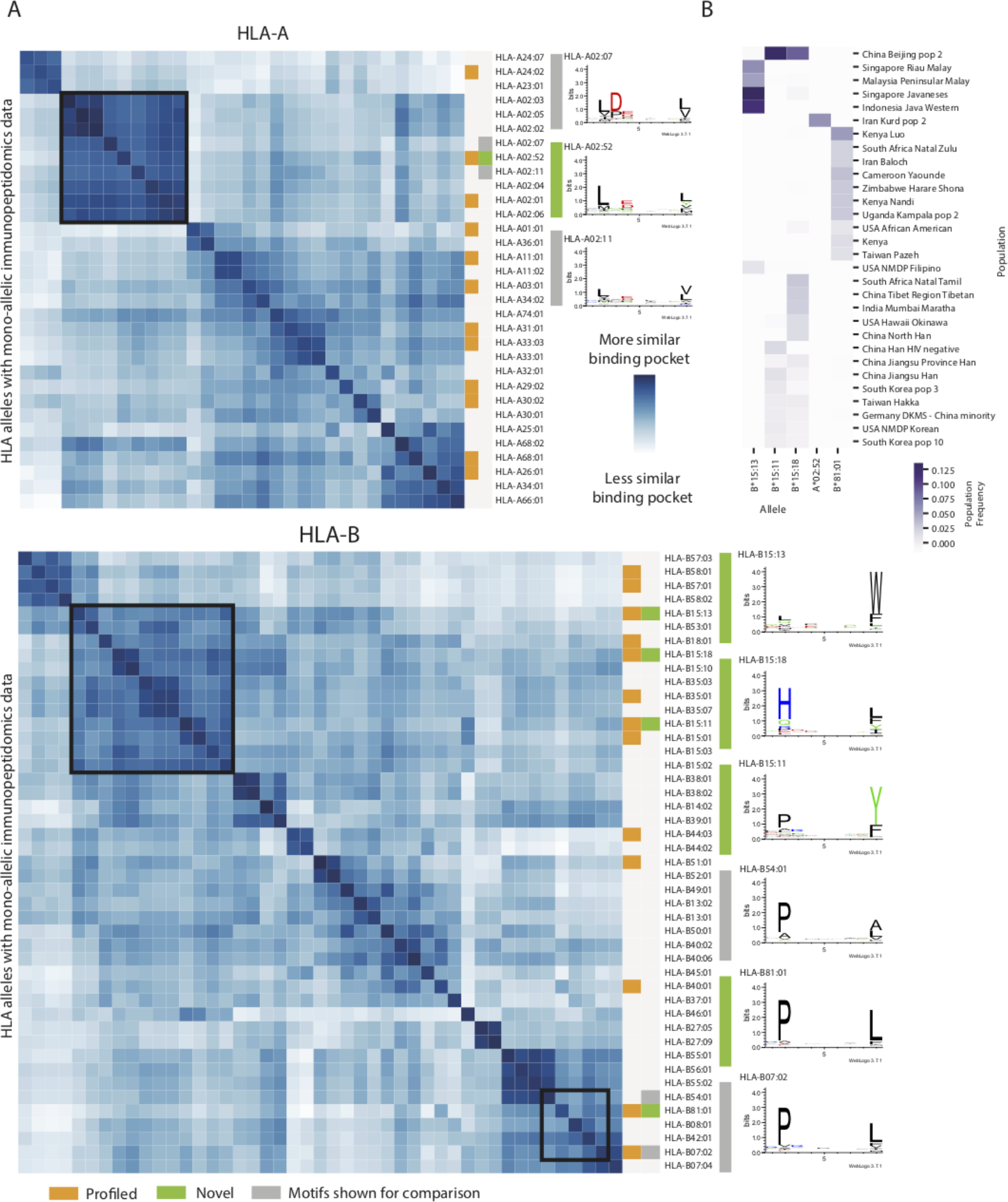
Binding pocket diversity and population frequencies of novel alleles. **(A)** Heatmaps for HLA-A and -B that represent the binding pocket similarity between alleles with mono-allelic immunopeptidomics data. Dark blue squares represent alleles that have very similar binding pockets while white squares represent alleles with divergent binding pockets. The 25 alleles profiled with our mono-allelic system are denoted in orange. The five alleles that have not previously been profiled are denoted in green. Motifs for these novel alleles are shown alongside motifs for related alleles in grey. Black boxes denote the cluster of alleles containing the newly profiled alleles. **(B)** A heatmap showing the frequencies of the five novel alleles in several populations of diverse world ethnicities. Dark purple denotes high population frequencies of the alleles and light purple denotes low population frequencies.

### Large scale data integration enhances the representativeness of HLA peptidome

Given the large amount of publicly available immunopeptidomics and binding affinity data from diverse tissue types and cell lines, we reasoned that a systematic integration of all available high-quality datasets would improve the representation of training data and enable feature engineering. Towards this end, we downloaded a large corpus of publicly available raw data (**Supp. Table 2**), comprising 2.15 million peptides from 512 experiments and covering 167 distinct alleles. A majority of the datasets are multi-allelic (384 experiments from 15 projects) (12,23,27,28,30–39), but there are also a large number of mono-allelic experiments (128 experiments from 6 projects) (7,13,40–44) and alleles with binding assay data (n=90 alleles) (26) (**Figure 3A**). As anticipated, the tissue types in the multi-allelic datasets have diverse expression profiles (**Figure 3B**). Furthermore, we observed significant differences in the gene expression profiles of transcripts from B721.221 cell lines (used in a large-scale mono-allelic dataset generated from B721.221 cell line with alleles from HLA-A, -B and -C) (7,13) and K562 cell lines (used in our in-house data generated) (**Figure 3C**). 21% of transcripts display significant differences in expression between B721.221 and K562 cell lines, and these variations impact key pathways that change the functionality of the cells (**Supp. Table 3**). Given the vast differences in underlying proteomes and antigen processing machinery of various tissue types, incorporation of these diverse data is particularly important for modeling presentation effectively.

**Figure 3:**
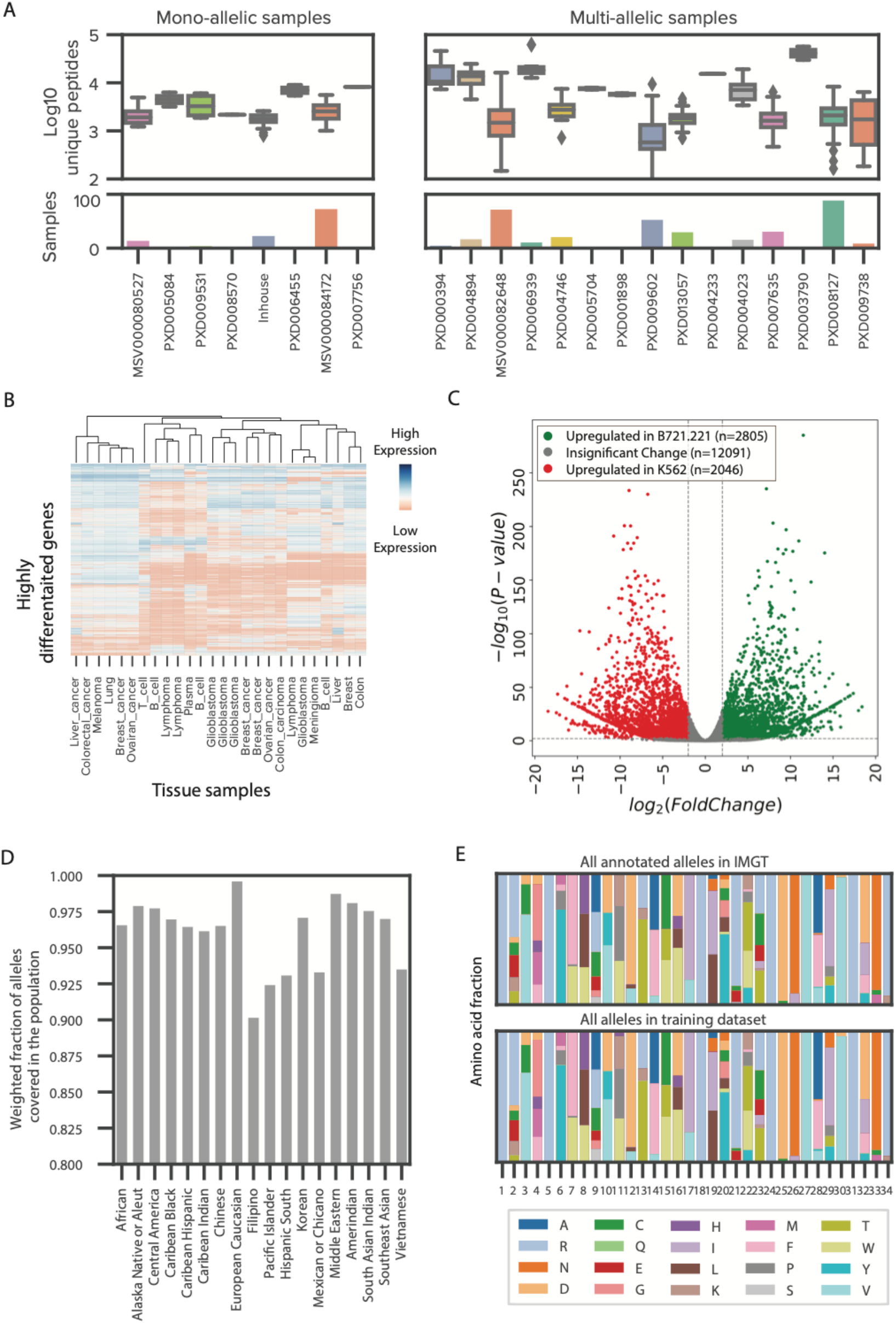
Systematic expansion of HLA ligandome through the incorporation of publicly available data. (A) Box plots representing the number of unique peptides per sample from mono-allelic and multiallelic immunopeptidomics samples that were reprocessed through our pipeline. Bar plot showing the number of samples for each project. Samples are colored according to their project. Peptide yields are log10 transformed. See Supplementary Table 2 for additional details. (B) A heatmap of expression values (TPM) of highly differentiated genes between tissue and tumor types of publicly available multi-allelic immunopeptidomics data. Low expression is shown with red, and high expression is shown with blue. (C) A volcano plot denoting the differential gene expression between the mono-allelic parental cell lines, B721.221 and K562. Gene transcripts with significant upregulation in B721.221 compared to K562 are shown in green while gene transcripts with significant upregulation in K562 compared to B721.221 are shown in red. Gene transcripts with no significant up or down regulation are shown in gray. (D) A bar plot denoting the weighted fraction of alleles in 18 ethnicity populations from the National Marrow Donor Program within the expanded training dataset, including mono-allelic cell lines profiled in house, public mono-allelic data, public multi-allelic data and binding assay data from IEDB. (E) Two stacked bar plots showing the frequencies of amino acids at each position in the pseudo binding pocket for all annotated alleles in IMGT (top) and all alleles from the expanded training dataset, including mono-allelic cell lines profiled in house, public mono-allelic data, public multi-allelic data and binding assay data from IEDB.

We evaluated the representation of the MHC alleles in our expanded dataset on two parameters -- population coverage and structural diversity of the binding pocket. We assessed population coverage using a simulation study that utilizes allele frequencies from the US National Marrow Donor Program as documented in the allele frequency net database (45). The alleles within our extended dataset represent an average coverage of 96% of observed alleles across various ethnic populations in the United States (**Figure 3D**). Our training data spans a wide range of allele binding pocket clusters that have vastly different motif characteristics, indicating a robust representation at the macro level. Since it is possible that relatively small differences in binding pocket sequences might lead to substantial differences in the presented motif, possibly due to the varied importance of amino acids in the binding pocket (46), we evaluated the representation at each position in the binding pocket. We observed a high degree of concordance in the distribution of frequencies between all known alleles and alleles in our expanded training dataset (**Figure 3E**). Of note, this concordance remains high even when we exclude alleles only present in multiallelic samples (**Supp. Figure 4A-B**). By expanding our in-house dataset using publicly available data, we not only enhance the breadth of our training data, both in terms of tissue-of-origin and allelic diversity, but also minimize potential biases introduced by data acquisition techniques, such as chromatography and peptide fragmentation approaches. Additionally, the expanded scale and scope helps minimize out-of-distribution modeling errors.

**Figure 4:**
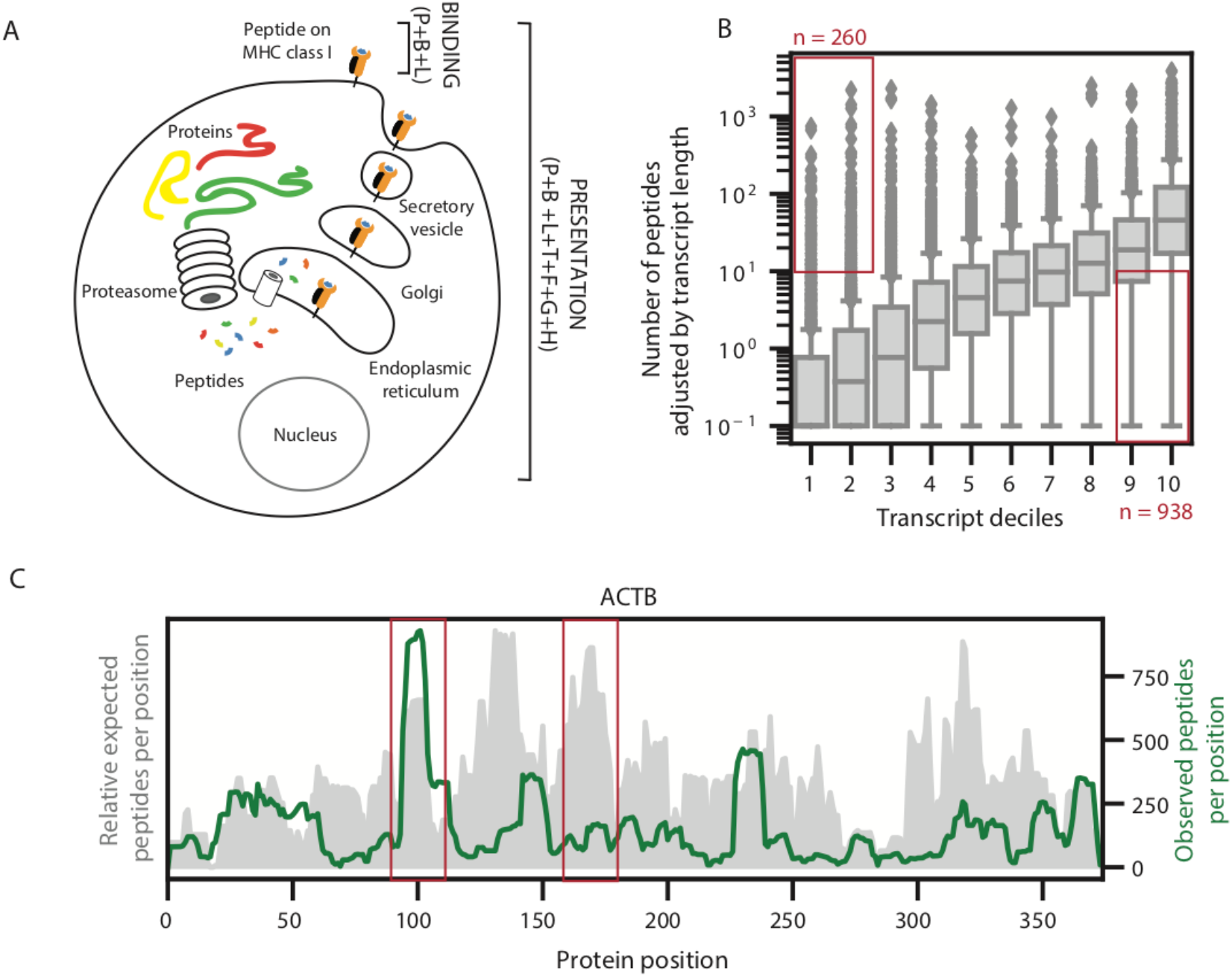
Modeling binding and presentation. **(A)** A schematic showing the difference between MHC binding and MHC presentation. MHC binding involves the ability of an MHC allele to bind to a paired peptide and is modeled with the peptide (P), allele binding pocket (B) and peptide length (L). MHC presentation involves all steps in the antigen processing pathway in addition to MHC binding and is modeled with the peptide (P), allele binding pocket (B), peptide length (L), gene expression (T), flanking regions around the peptide (F), propensity of the gene to engender peptides (G) and propensity of the region within the gene to engender peptides (H). **(B)** Boxplots representing the distribution of peptides per transcript observed in the re-processed multi-allelic immunopeptidomics data across transcript deciles. The peptides observed are normalized by transcript length. Red boxes denote the transcripts that generate many observed peptides despite low expression levels and transcripts that generate few observed peptides despite high expression levels. **(C)** The distributions of expected and observed peptides from across the ACTB protein. Expected peptides, shown in grey, are generated by summing the number of frequent alleles predicted to bind each peptide (Rank < 2 by netMHCpan4.0). The 30 most frequent alleles in the re-processed multiallelic immunopeptidomics dataset were used for the analysis. Observed peptides are measured from the re-processed multi-allelic immunopeptidomics data and are shown in green.

### Modeling peptide-MHC presentation using novel features encapsulating antigen processing

The unbiased and ex-vivo nature of mass-spectrometry-based immunopeptidomics data uniquely lends itself to model both MHC-peptide binding and presentation. Modeling *binding* entails learning the motif preferences of HLA alleles and their cognate peptides, whereas modeling *presentation* includes learning propensities of upstream antigen processing in addition to binding motifs (**Figure 4A**). Several features that capture various aspects of antigen processing and the biophysical characteristics of MHC-peptide binding have been discussed in the literature, the most prominent of these are peptide sequence, binding pocket sequence, expression of source protein, cleavage patterns and various propensities for presentation that are not captured by expression (12). We use a combination of widely applied and novel features to model MHC-peptide binding and presentation.

We modeled binding using three features: the amino acid sequence of the peptide ligand (P), the binding pocket pseudo sequence of the presenting allele (B) and peptide length (L). In addition to these three binding-specific features, we integrated four additional features into our presentation model that encapsulate antigen processing. First, as evidence from our mono-allelic immunopeptidomics data, the abundance of the source protein has a strong influence on peptide presentation (**Figure 1D**). Accordingly, we used a transformed value of transcript expression (T) as a surrogate for protein expression. Second, our data also showed the strong preference of amino acids flanking the presented peptides for proteasomal cleavage; therefore, we incorporated the five amino acids upstream and downstream of each peptide (F) as an additional feature (**Figure 1E**). It has been discussed previously that certain proteins or regions within proteins have a higher propensity for MHC presentation (23,47,48). We have noticed similar trends in the large set of public data that we had downloaded and reprocessed systematically (**Supp. Table 2**). We evaluated the dependence of the number of peptides presented (adjusted for transcript length) as a function of transcript expression of its cognate gene. As expected, we observed a positive correlation in the median values, but we also observed several outliers that do not follow this trend (data points highlighted in red boxes, **Figure 4B**). We modeled this skew (propensity) and developed a gene propensity score (G) as our third feature (**Supp. Figure 5A**, see Methods). Similarly, we noticed positional preferences within proteins when we compared observed and expected peptides from various locations within a protein. An anecdotal example of such a skew in Actin Beta gene (ACTB) is shown in **Figure 4C**. Briefly, position specific coverage of observed peptides was contrasted with expected coverage based on predictions for the thirty most frequent alleles in our dataset. Importantly, we found a lack of correlation between predicted and observed peptide profiles across the entire proteome (**Supp. Figure 5B**). We modeled this phenomenon using position specific coverage of immunopeptides in our large dataset and developed a hotspot score (H) as our final feature (see Methods). An ensemble of these propensity scores and other features described above facilitate the modeling of antigen processing and surface presentation.

### Systematically incorporating binding affinity, mono-allelic and multi-allelic data into a composite model

Both in vitro binding assays and mono-allelic immunopeptidomics experiments produce allelespecific data, but assigning a peptide to its cognate allele, also known as deconvolution, is a key challenge in modeling multi-allelic data. Several approaches have been applied to perform deconvolution (10–12). We utilized our comprehensive binding affinity and mono-allelic data sets to develop a model-based deconvolution approach to convert multi-allelic data into pseudo monoallelic data (**Figure 5A, Supp. Table 4**). First, we generated a binding prediction model using exclusively mono-allelic data (in-house mono-allelic data, public mono-allelic data and in-vitro binding assay data extracted from IEDB). Unlike the binding affinity data from IEDB that generates both positive and negative examples, immunopeptidomics data only generates positive examples, so we generated negative examples using a large set of random peptides from the human proteome at a 1:20 ratio. A large proportion of negative examples helps minimize the risk of learning random or systematic biases in synthetically generated negative examples. This monoallelic model (MONO-Binding) is able to accurately predict motifs for alleles excluded from the training dataset (**Figure 5B, Supp. Figure 6**). Given our confidence in the model trained with allele-specific data (MONO-Binding), we applied it to each of the multi-allelic datasets to assign each peptide in a sample to its cognate allele and generate pseudo mono-allelic data. From 289 multi-allelic sources representing 118 unique HLA alleles, we began with over 1.15 million peptides and successfully mapped about 700,000 peptides to alleles (~60% mapping rate). Then, the mono-allelic and pseudo mono-allelic data were merged to train a comprehensive binding model (SHERPA-Binding). Finally, SHERPA-Binding was used as a primary model (p-model) or feature for our SHERPA-Presentation model. This final model incorporates all presentation features but is trained exclusively on mono-allelic data to avoid overfitting to the multi-allelic datasets that generated the gene propensity and hotspot features. As expected, when we examined the feature importance for each of the models, we observe the peptide anchor residues and diverse binding pocket amino acids have the greatest influence over the SHERPA-Binding model, and SHERPA-Binding has the greatest influence over the SHERPA-Presentation model (**Supp. Figure 7**). For all models, prediction probabilities were calibrated into percentile ranks to remove allele-specific biases as described earlier (15).

**Figure 5:**
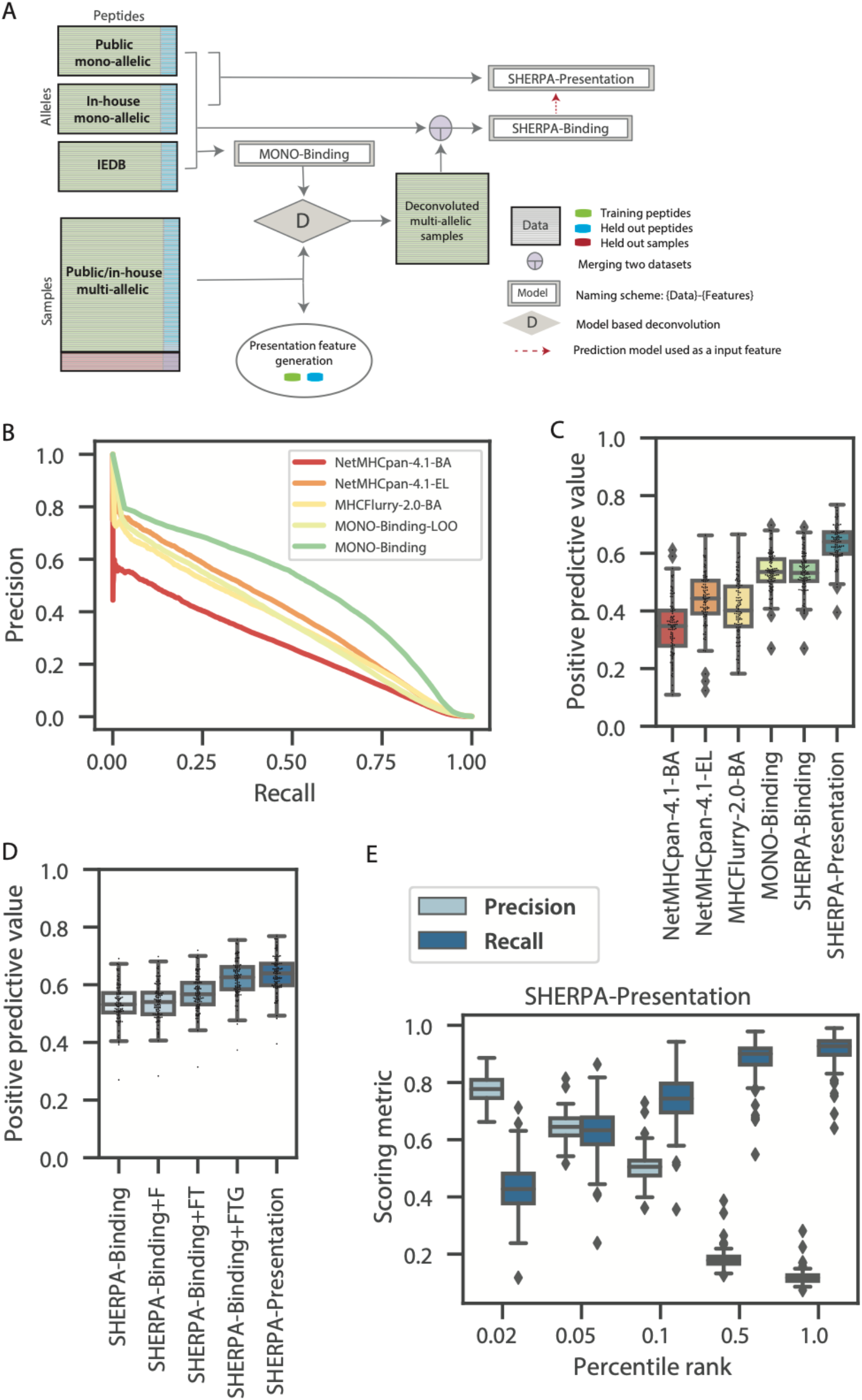
Overview of composite modeling approach and model performance. **(A)** A schematic of the composite modeling approach. Inhouse mono-allelic immunopeptidomics data, public mono-allelic immunopeptidomics data and IEDB data are used to train MONO-Binding. MONO-Binding is used to deconvolute the multi-allelic immunopeptidomics data to create pseudo mono-allelic data. All mono-allelic and pseudo-mono-allelic data is combined to train the SHERPA-Binding model. The SHERPA-Binding model is used as a feature along with other presentation features to train the SHERPA-Presentation model on mono-allelic immunopeptidomics data. **(B)** A precision-recall curve demonstrating the predicted pan-performance on unseen alleles (MONO-Binding-LOO) compared to MONO-Binding and NetMHCpan4.1-BA, NetMHCpan-4.1-EL, MHCFlurry-2.0-BA. A model was trained for each allele with the data for that allele excluded from the training dataset. The MONO-Binding-LOO curve represents the predictions from each of the models on the test data of the allele excluded from the training data. **(C-D)** Boxplots denoting the distributions of positive predictive values (top 0.1%) across alleles within the mono-allelic immunopeptidomics held-out test data. Distributions are shown for **(C)** NetMHCpan4.1-BA, NetMHCpan-4.1-EL, MHCFlurry-2.0-BA, MONO-Binding, SHERPA-Binding and SHERPA-Presentation and **(D)** SHERPA-Binding, SHERPA-Binding+F, SHERPA-Binding+FT, SHERPA-Binding+TTG and SHERPA-Presentation. **(E)** Boxplots showing the distribution of precision and recall values across alleles in the mono-allelic immunopeptidomics data for SHERPA-Presentation across several percentile rank thresholds. A percentile rank of 0.1 is selected as the optimal threshold.

### Benchmarking the performance of SHERPA binding and presentation models

To evaluate our models, we first tested their performance on ~10% held-out mono-allelic data (positive examples), mixed with negative examples in a 1:999 ratio to mimic true ratio (5,23–26). We chose this prevalence to reflect the underlying characteristics of peptide presentation (5,23–26). The MONO-Binding model significantly outperformed NetMHCpan-4.1-BA, NetMHCpan-4.1-EL and MHCFlurry-2.0-BA (0.53, 0.34, 0.44 and 0.40, respectively, **Figure 5C, Supp. Data 1A**). When we restricted the input data to be only the publicly available mono-allelic data (PUBLIC-Binding), SHERPA still outperformed the other three models, suggesting that the XGBoost modeling approach or our strict data curation added significant value (**Supp. Figure 8A**). The binding model trained on the full dataset (SHERPA-Binding), comprising both mono-allelic and pseudo mono-allelic data, had the same positive predictive value (PPV) compared to MONO-Binding, a model trained on mono-allelic data alone (0.53 and 0.53, respectively; **Figure 5C**). SHERPA-Presentation has a better PPV compared to SHERPA-Binding, attesting to the utility of presentation-specific features. We further evaluated the contribution of each of our features and confirmed that each feature has an additive effect on performance (**Figure 5D**). Flanking regions had a relatively minor impact on performance (~0.01 PPV gain compared to SHERPA-Binding). We found the largest performance gain with the addition of abundance (TPM, ~0.03 increase in PPV above SHERPA-Binding+F) and gene propensity (~0.05 increase in PPV above SHERPA-Binding+FT), but we also observed significant gain with the addition of the hotspot score (~0.01 PPV above SHERPA-Binding+FTG). For an orthogonal allele-specific validation, we assessed the performance of the models using the held out IEDB data. Of the comparison models, NetMHCpan-4.1-BA had the best performance. MONO-Binding and SHERPA-Binding performed similarly to NetMHCpan-4.1-EL and MHCFlurry-2.0-BA (**Supp. Figure 8B, Supp. Data 1B**). Finally, we evaluated the precision and recall at various rank values for both our multi-allelic binding and presentation models. We observed a consistently and significantly higher precision and recall compared to other models at all rank values (**Figure 5E, Supp. Figure 8C-E**). Using this analysis, we empirically determined a rank threshold of 0.1 to be our definition of a ‘binding’ or ‘presented’ peptide.

### Experimental validation of SHERPA using tissue samples

To understand the utility of our prediction models in a patient sample, we performed immunopeptidomics on tumor samples from seven lung cancer patients and five colorectal cancer patients. In these experiments, we observed strong peptide yields (4043 median, **Supp. Figure 9A, Supp. Table 5, Supp. Data 1C**). As expected, we did not observe any neoantigens from these patients in the immunopeptidomics data, as data dependent acquisition (DDA) based IP-MS protocols lack the required sensitivity of detection. A total of 46 alleles were represented across the 12 patients, and two of the alleles were outside of our training dataset. Thus, testing our performance on these samples provided an opportunity to assess our prediction accuracy on trained alleles, as well as the pan-allelic prediction capability on untrained alleles. To establish a prediction score for each peptide across all six potential patient alleles (patient-centric rank), we used the rank from the allele with the lowest rank to represent how well a peptide is bound or presented by that patient. We observed a strong performance of our models, which recovered 1.15 times more experimentally observed peptides among those that were predicted to be bound or presented than the next best model (MHCFlurry-2.0-BA) (**Figure 6A**). Further, the multi-allelic models outperformed the exclusively allele-specific models, highlighting the importance of the multi-allelic deconvolution approach. Of note, we observed one sample with significantly lower recall than those from the other patients across all models. We explored that patient further and found that the patient has human leukocyte antigen (HLA) loss of heterozygosity across two HLA genes (-A and -B; **Supp. Figure 9B**), highlighting the importance of considering a comprehensive view of the patient HLA features during neoantigen prediction. Although all models used a percentile ranking approach, we found that MHCFlurry-2.0-BA predicts an average of 30 percentile rank for negative peptides (compared to the expected ~50 percentile rank for all other models), suggesting that the sensitivity metric used in this analysis may be overrepresenting MHCFlurry's performance (**Supp. Figure 9C-D**). To further corroborate our results, we performed the same analysis in the ~10% of the peptides from tissue data from ovarian cancer and colorectal cancer studies that had been held-out from multi-allelic training (27, 28). We found similar trends across both of these external datasets, on data that were not part of our training data, with the SHERPA-presentation model showing the best results, showing 1.03-1.22 times more experimentally observed peptides predicted than MHCFlurry-2.0-BA (**Figure 6B-C**).

**Figure 6:**
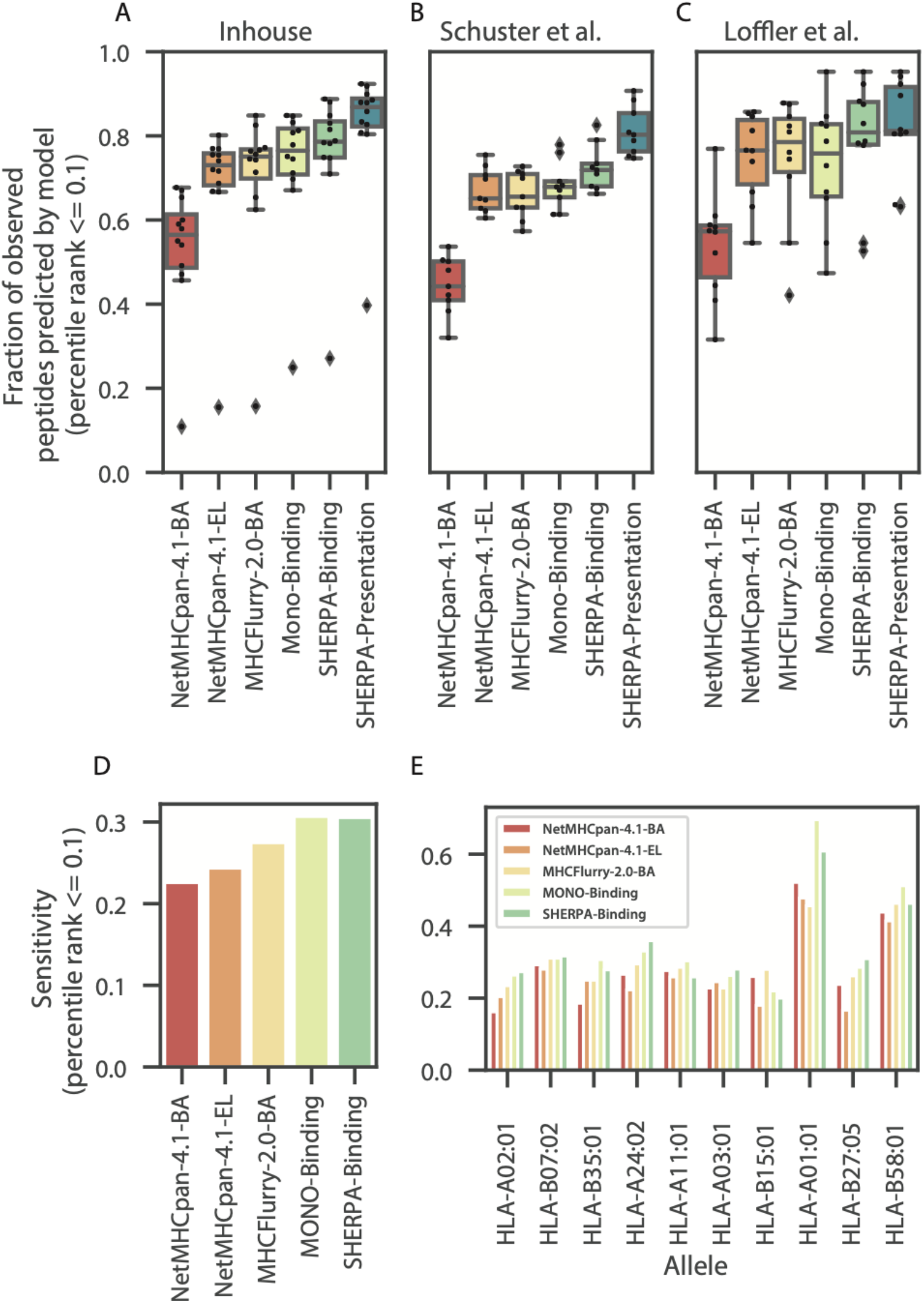
Performance of SHERPA on tissue samples and immunogenic epitopes. Boxplots showing the distribution of prediction performance across **(A)** tumors profiled with immunopeptidomics in-house (lung and colorectal, left), **(B)** by Schuster et al. (ovarian, middle) and **(C)** Loffler et al. (colorectal, right). Performance is defined as the fraction of peptides observed with immunopeptidomics that are predicted to bind in the top 0.1% of all peptides percentile rank ≤ 0.1). Performance is shown for the following models: NetMHCpan4.1-BA, NetMHCpan-4.1-EL, MHCFlurry-2.0-BA, MONO-Binding, SHERPA-Binding and SHERPA-Presentation. **(D-E)** Bar plots showing the sensitivity of NetMHCpan4.1-BA, NetMHCpan-4.1-EL, MHCFlurry-2.0-BA, MONO-Binding and SHERPA-Binding on the Chowell et al. immunogenicity dataset: **(D)** performance across all epitopes and **(E)** performance across high frequency alleles.

### Immunogenicity

Although the SHERPA models do not specifically predict the likelihood that a peptide will be immunogenic, MHC presentation is a gatekeeping step of an immunogenic response. Thus, one would expect that all immunogenic peptides are successfully bound and presented on the cell surface. As an additional evaluation, we used the dataset of immunogenic peptides described by Chowell et al to evaluate the ability of the models to predict these epitopes (29). Across the entire dataset, we observe the best performance by the MONO-Binding model and the SHERPA-Binding model (**Figure 7D, Supp. Data 1D**). We see variable but similar performance across the specific alleles. Of note, we observe a 1.11 fold improvement over the next best model for HLA-A*02:01, the most studied allele (**Figure 7E**). Together, this evidence suggests that the performance of the SHERPA models is generalizable and resilient to data sources.

## Discussion

Many personalized cancer immunotherapies require precise neoantigen identification. Although advances in next generation sequencing technologies have enabled large-scale survey of putative neoantigens, algorithmic neoantigen ranking continues to have limited accuracy. Our work seeks to improve precision neoantigen discovery by addressing both the training data and modeling approach. Since prediction models are highly dependent upon their training data, we generated immunopeptidomics data using mono-allelic cell lines that unambiguously map a peptide to an HLA allele. To ensure generalizability of SHERPA beyond our model system, we further expanded the HLA-peptidome by incorporating publicly available mono-allelic immunopeptidomics data from different parental cell lines, allele-specific binding array data and multi-allelic tissue immunopeptidomics data. Moreover, we developed a modeling approach that integrates the data types strategically and mines the breadth of data to model antigen processing across a variety of tissue types. It also accounts for real world data by accounting for diploid allele data.

While a combination of several factors leads to marked improvements in performance of SHERPA, we believe that three aspects set it apart. First, SHERPA employs the largest training data of any neoantigen prediction tool to our knowledge. Peptides from 167 unique human alleles and antigen processing from 30 unique expression profile backgrounds ensure the representativeness of our training data. Second, we designed features that mine extensive tissue-based immunopeptidomics data to model presentation and demonstrate ~0.06 improvement in our mono-allelic PPV assessment with their inclusion. Third, SHERPA's composite modeling approach for integrating multiple data types reduces inherent biases caused by a single data source and allows for robust multi-allelic deconvolution.

Our work is timely because it addresses two critical needs in the field. First, the Human Immuno-Peptidome Project outlined the need for large cohort immunopeptidomics studies in their effort to comprehensively map the human immuno-peptidome (49). With this project, we contribute a new set of mono-allelic immunopeptidomics data on a different cell line background than previous large studies and a previous set of lung and colorectal tumor tissues. Second, the scientific community has an urgent need for representing diverse ethnic populations in medical research (18). We addressed this issue by profiling five novel HLA alleles that are frequent in underrepresented Asian, African and Middle Eastern populations.

Though our work shows great improvement, both SHERPA and the field of neoantigen prediction continues to face some key challenges. We integrated different data types (binding array and immunopeptidomics) to decrease bias and increase generalizability, whereas the vast majority of previous pMHC data was produced using very similar protocols. Accordingly, developing newer approaches to affinity purification other than the commonly used W6/32 antibody immunoprecipitation, applying newer data acquisition strategies such as data independent acquisition, and generating immunopeptidomics data under different cell states are avenues for future improvement. In addition, gene expression is an imperfect surrogate for protein abundance. Other methods, such as ribosome footprinting, may provide a better estimate of protein levels. Moreover, some of the features used in the SHERPA-Presentation model are exclusively designed to the human proteome without holding out specific proteins or protein sub-regions, so the results of the SHERPA-Presentation model should be interpreted in light of the inclusion of all proteins and protein-regions in the feature generation process. The SHERPA-Binding-FT model is a more accurate representation of performance on non-human candidates. Finally, SHERPA does not specifically predict the likelihood that a ligand will be immunogenic; however, SHERPA captures a higher percentage of immunogenic epitopes than other methods, suggesting that peptides predicted by SHERPA will be more likely to be immunogenic.

In summary, we demonstrate that SHERPA is an important method that balances the tension between the clarity of mono-allelic motifs and the generalizability of tissue data while consistently outperforming previous models. Its improved performance is expected to be useful in neoantigen prediction for immunotherapy.

## Data Availability

The mass spectrometry proteomics data have been deposited to the ProteomeXchange Consortium via the PRIDE partner repository with the dataset identifier PXD023064 (50). Evaluation data can be downloaded with the following link: https://drive.google.com/drive/folders/1y3IBELIU5TgUaEiWrPANxRlTVUhYQGGp?usp=sharing.

## Conflict of Interest

R.M.P., D.M., S.D., C.A., S.V.Z., N.P., J.H., G.B., S.D., R.M., J.W., R.C. and S.M.B are full time employees of Personalis. M.P.S. co-founded Personalis. Personalis Inc provided the funding for this project.

## Acknowledgements

The cell line transfections were conducted at Thermo Fisher Scientific. The immunopeptidomics experiments were conducted at Cayman Chemicals and MS Bioworks, LLC. We thank Josette Northcott for her help in interpretation of the flow cytometry data. We thank Eric Levy, Euan Ashley, Russ B. Altman and Atul Butte for their helpful feedback on the manuscript.

## Abbreviations

MHC: major histocompatibility complex
pMHC: major histocompatibility complex-peptide
HLA: human leukocyte antigen
LC-MS/MS: liquid chromatography with tandem mass spectrometry
TPM: transcripts per million
FDR: false discovery rate
ELISA: enzyme-linked immunosorbent assay
GFP: green fluorescent protein
NMDP: National Marrow Donor Program
ATCC: American Type Culture Collection
IMGT: international ImMunoGeneTics information system
IEDB: Immune Epitope Database and Analysis Resource
LOO: Leave one out model
SHERPA: Systematic HLA Epitope Ranking Pan Algorithm
P-models: primary models
B: binding pocket (model feature)
P: peptide (model feature)
L: peptide length (model feature)
T: protein abundance as measured by TPM (model feature)
F: flanking regions (model feature)
G: gene propensity (model feature)
H: hotspot score (model feature)

## Supplemental Data

This article contains supplemental data (**Supp. Table 1-5, Supp. Data, Supp. Code, Supp. Figures 1-9**).

