## Supplemental Figures and Legends for "Precision neoantigen discovery using large-scale immunopeptidomes and composite modeling of MHC peptide presentation"

**This file includes:**

Supplementary Table Legends 1-5
Supplementary Data Legends 1
Supplementary Code Legends
Supplementary Figures 1-9

### Supplementary Tables

**Supplementary Table 1: Peptide yields of mono-allelic immunopeptidomics data generation.** (A) The number of unique peptides measured from each of the 25 mono-allelic immunopeptidomics experiments. (B) The unique peptides identified for each allele.

**Supplementary Table 2: Public data incorporated into the expanded dataset.** (A) The project, sample/allele and category (single allele or multi-allelic) used in the study. (B) The set of processed peptides used from IEDB.

**Supplementary Table 3: Enriched pathways between popular mono-allelic parental cell lines.** The gene ontology (GO) pathways that are (A) enriched or (B) depleted in K562 compared to B721.221. P-values and FDRs are also shown.

**Supplementary Table 4: Overview of models.** A table containing all of the details about each model: model name, datasets used for training, number of alleles derived from mono-allele datasets, number of alleles derived from multi-allele datasets, number of total alleles and features.

**Supplementary Table 5: Overview of tissue immunopeptidomics samples.** (A) The number of unique peptides derived from the immunopeptidomics experiment of each tumor sample. (B) The HLA types of the patients. (C) The unique peptides identified for each patient.

### Supplementary Data

**Supplementary Data 1: Model predictions on test data.** (A) Mono-allelic mass spectrometry data. (B) IEDB data. (C) Multi-allelic tumor tissue data. (D) Immunogenic epitope data. Data can be downloaded with the following link: https://drive.google.com/drive/folders/1y3IBELIU5TgUaEiWrPANxRITVUhYQGGp?usp=sharing.

### Supplementary Code

**Peaks_post_processing.py -** Script to process the PEAKS output to generate peptide lists.

### Supplementary Figures

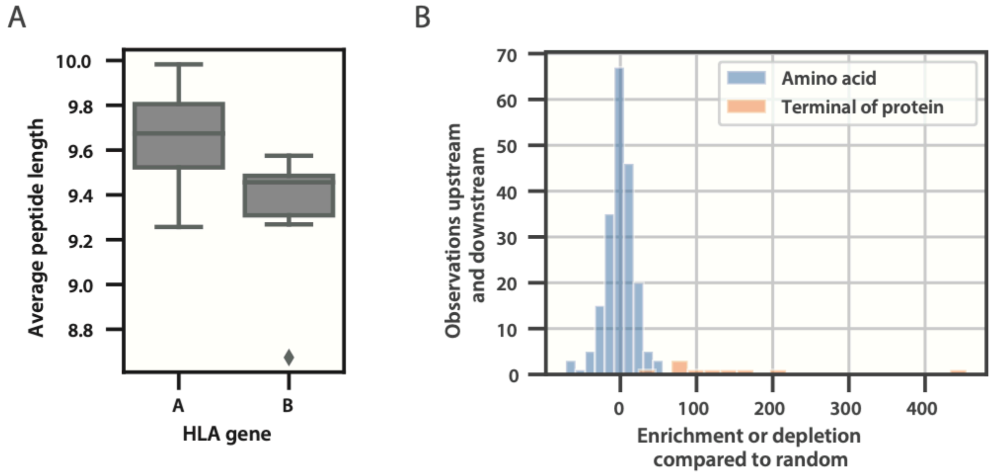


**Supplementary Figure 1:** Generation of mono-allelic immunopeptidomics data. (A) A box plot showing the difference in average peptide length for HLA-A alleles and HLA-B alleles. Two-sided t-test, p=0.006. (B) A histogram comparing the distribution of enrichment or depletion of amino acids upstream and downstream of the peptide of interest. The blue distribution shows all amino acids. The orange distribution shows the two termini of the protein. Two-sided t-test, p=3.77e-32.

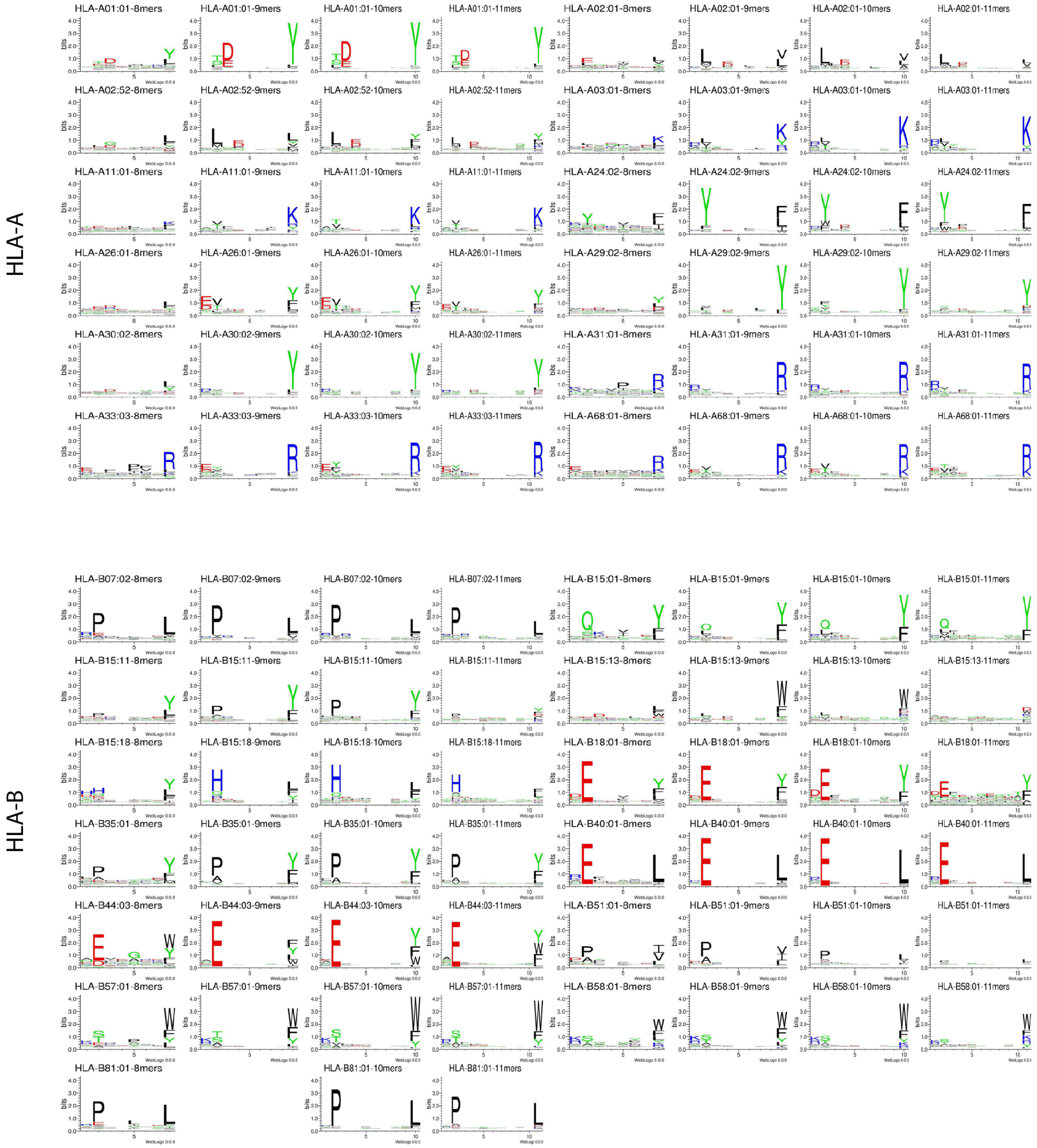


**Supplementary Figure 2:** Motifs of peptides from immunopeptidomics of 25 mono-allelic cell lines. Motifs are shown for HLA-A and -B alleles for peptides of length 8, 9, 10 and 11 amino acids.

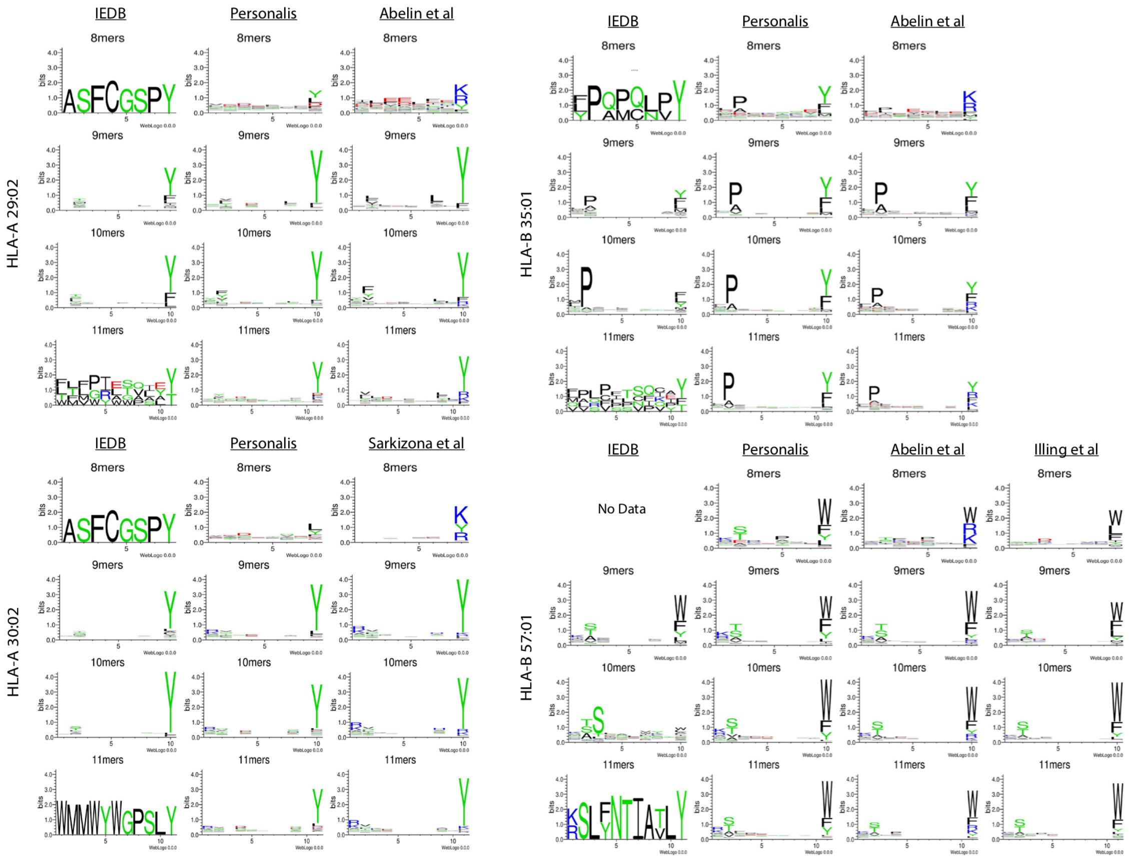


**Supplementary Figure 3:** Motifs of peptides for alleles with tryptic digest signatures in public datasets. Motifs for peptides of length 8, 9, 10 and 11 are shown for HLA-A29:02, HLA-A30:02, HLA-B35:01 and HLA-B57:01. Motifs from peptides derived from IEDB, in house mono-allelic cell lines and public mono-allelic cell lines are shown for comparison.

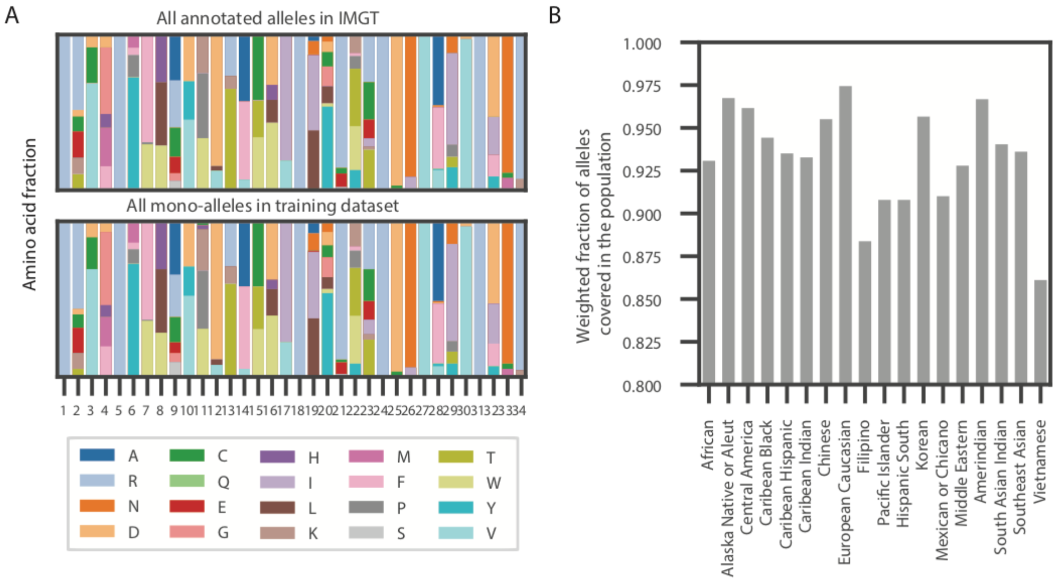


**Supplementary Figure 4:** Binding pocket representation and population frequency coverage of alleles in mono-allelic data. **(A)** Two stacked bar plots showing the frequencies of amino acids at each position in the pseudo binding pocket for all annotated alleles in IMGT (top) and all alleles from the expanded training dataset, including mono-allelic cell lines profiled in house, public mono-allelic data and binding assay data from IEDB. **(B)** A bar plot denoting the weighted fraction of alleles in 18 ethnicity populations from the National Marrow Donor Program within the expanded training dataset, including mono-allelic cell lines profiled in house, public mono-allelic data and binding assay data from IEDB.

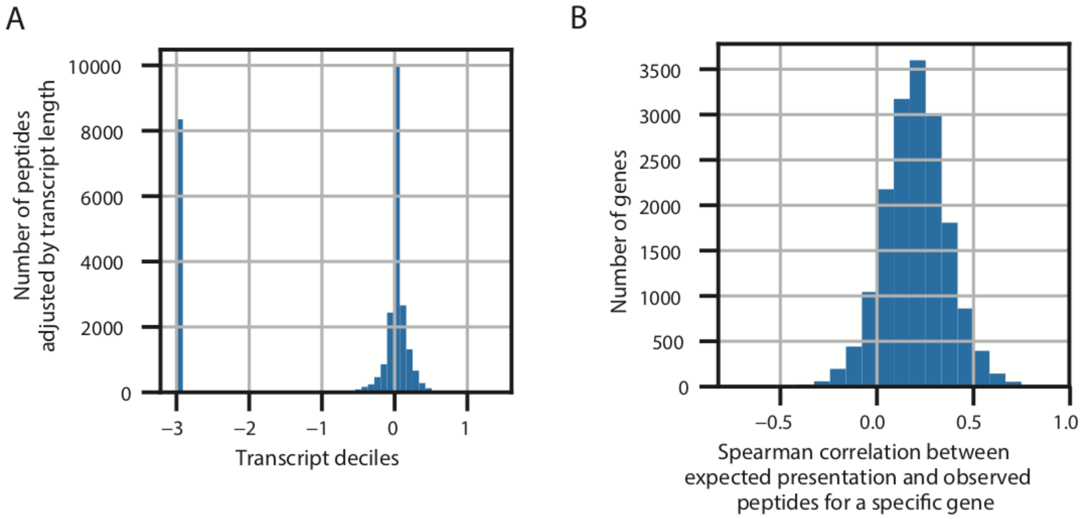


**Supplementary Figure 5:** Gene and within gene propensity feature overview. **(A)** The distribution of the gene propensity feature across transcripts. The feature is defined as the log10 transformation of the number of observed peptides for a transcript in the multi-allelic immunopeptidomics data divided by the expected number of peptides (as defined as the average TPM multiplied by the gene length). Transcripts without any observed expression across all samples were assumed to be pseudo genes and were automatically assigned a value of -3. **(B)** The distribution of the correlations between the number of observed peptides and the number of expected peptides (as measured through prediction) at each peptide within a protein across all individual genes.

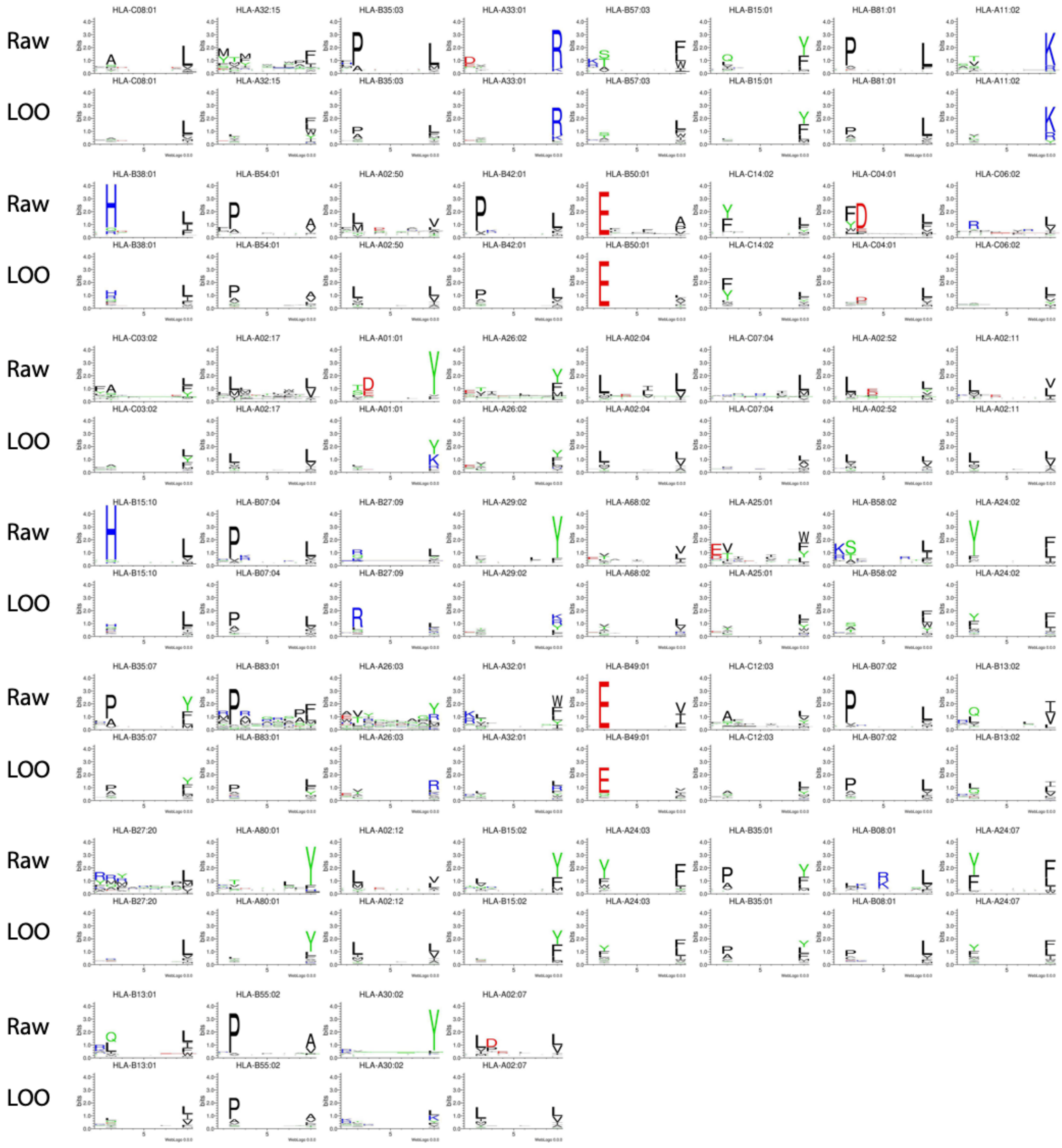
Raw
LOO
HLA-C08:01
HLA-A32:15
HLA-B35:03
HLA-A33:01
HLA-B57:03
HLA-B15:01
HLA-B81:01
HLA-A11:02
HLA-B38:01
HLA-B54:01
HLA-A02:50
HLA-B42:01
HLA-B50:01
HLA-C14:02
HLA-C04:01
HLA-C06:02
HLA-C03:02
HLA-A02:17
HLA-A01:01
HLA-A26:02
HLA-A02:04
HLA-C07:04
HLA-A02:52
HLA-A02:11
HLA-B15:10
HLA-B07:04
HLA-B27:09
HLA-A29:02
HLA-A68:02
HLA-A25:01
HLA-B58:02
HLA-A24:02
HLA-B35:07
HLA-B83:01
HLA-A26:03
HLA-A32:01
HLA-B49:01
HLA-C12:03
HLA-B07:02
HLA-B13:02
HLA-B27:20
HLA-A80:01
HLA-A02:12
HLA-B15:02
HLA-A24:03
HLA-B35:01
HLA-B08:01
HLA-A24:07
HLA-B13:01
HLA-B55:02
HLA-A30:02
HLA-A02:07

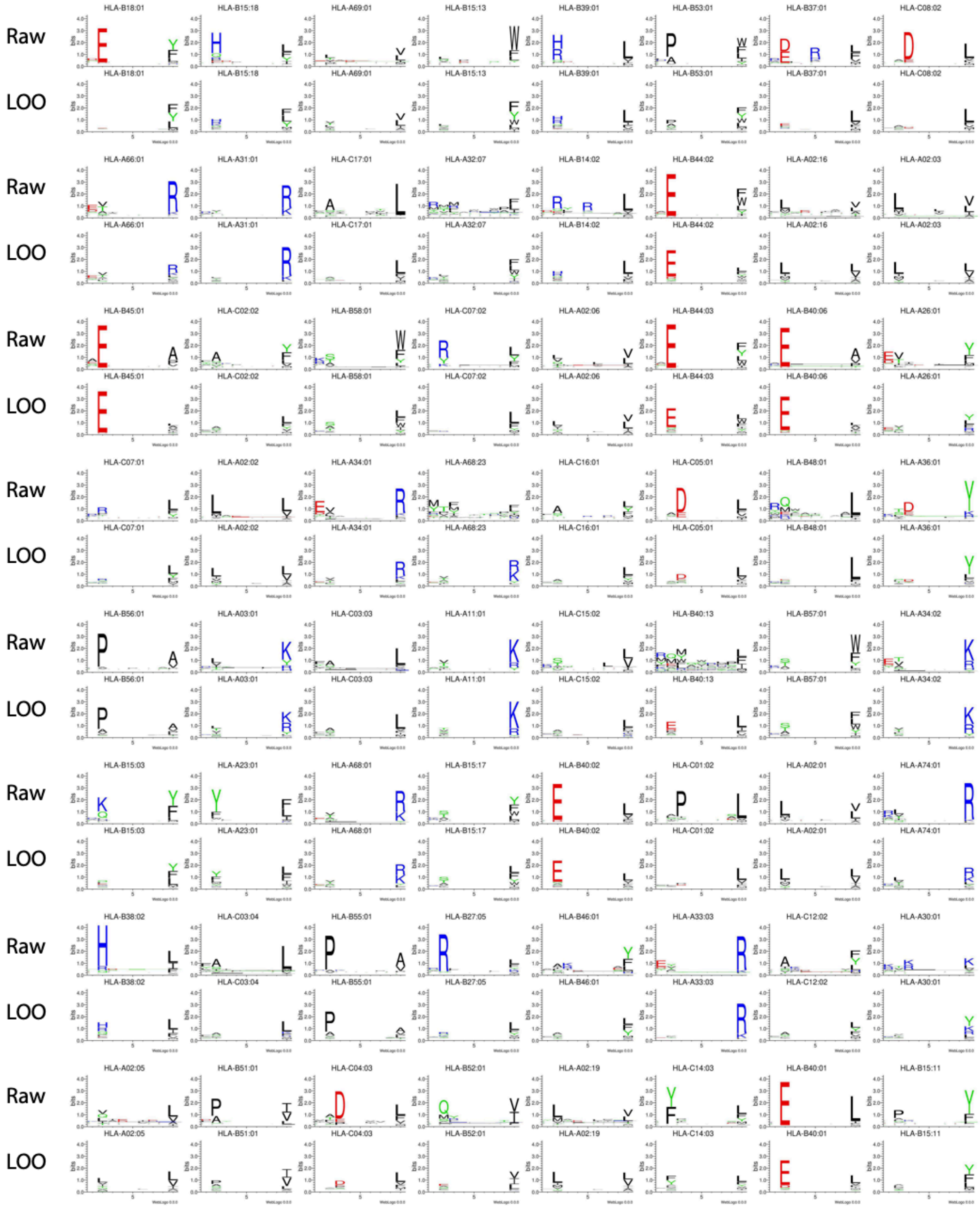


**Supplementary Figure 6: Comparison between raw immunopeptidomics motifs and pan-allelic predictions.** For all alleles with at least 50 peptides in the full mono-allelic dataset (in-house, public and IEDB), two motifs are shown. First, the ‘raw’ motif derived from observed peptides for that allele from the full mono-allelic dataset is shown on top. Second, the ‘LOO’ motif derived from the peptides predicted to bind to the allele by a model trained without the allele (MONO-LOO) from a set of random peptides.

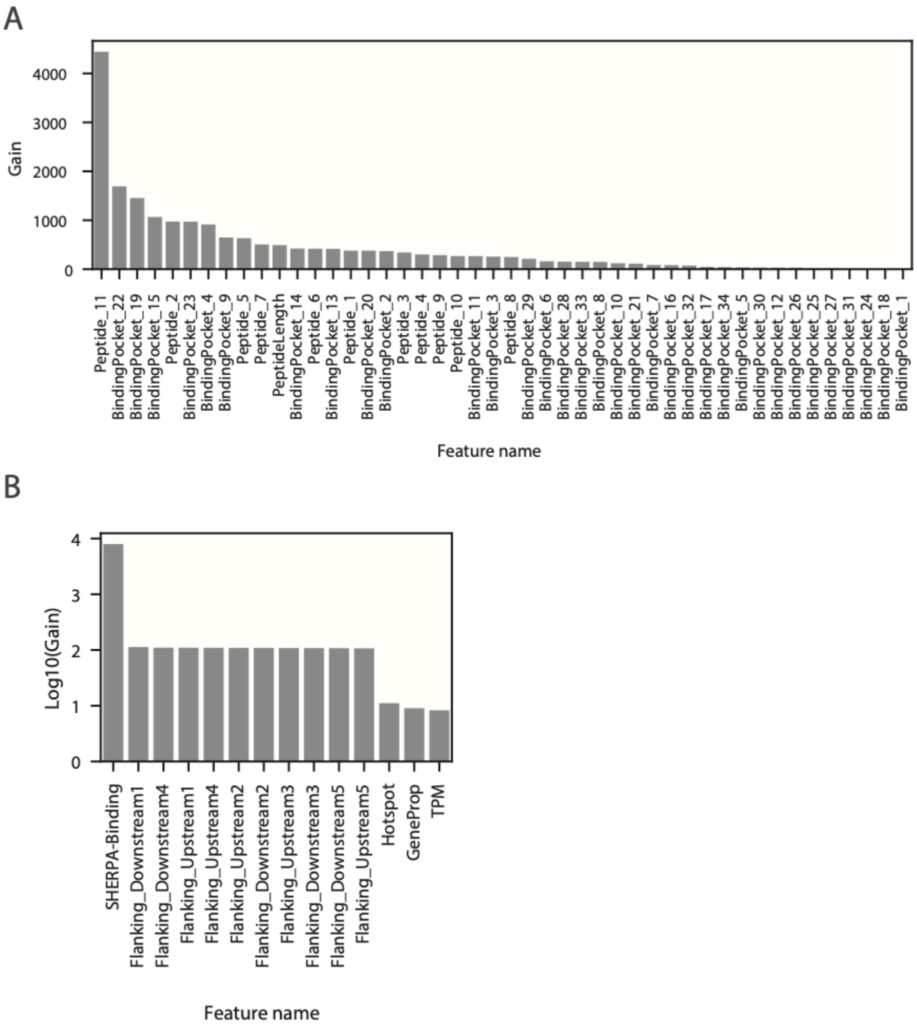


**Supplementary Figure 7: Overview of model feature importance. (A-B)** Bar plots denoting the feature importance (shown as 'gain') of **(A)** SHERPA-Binding and **(B)** SHERPA-Presentation.

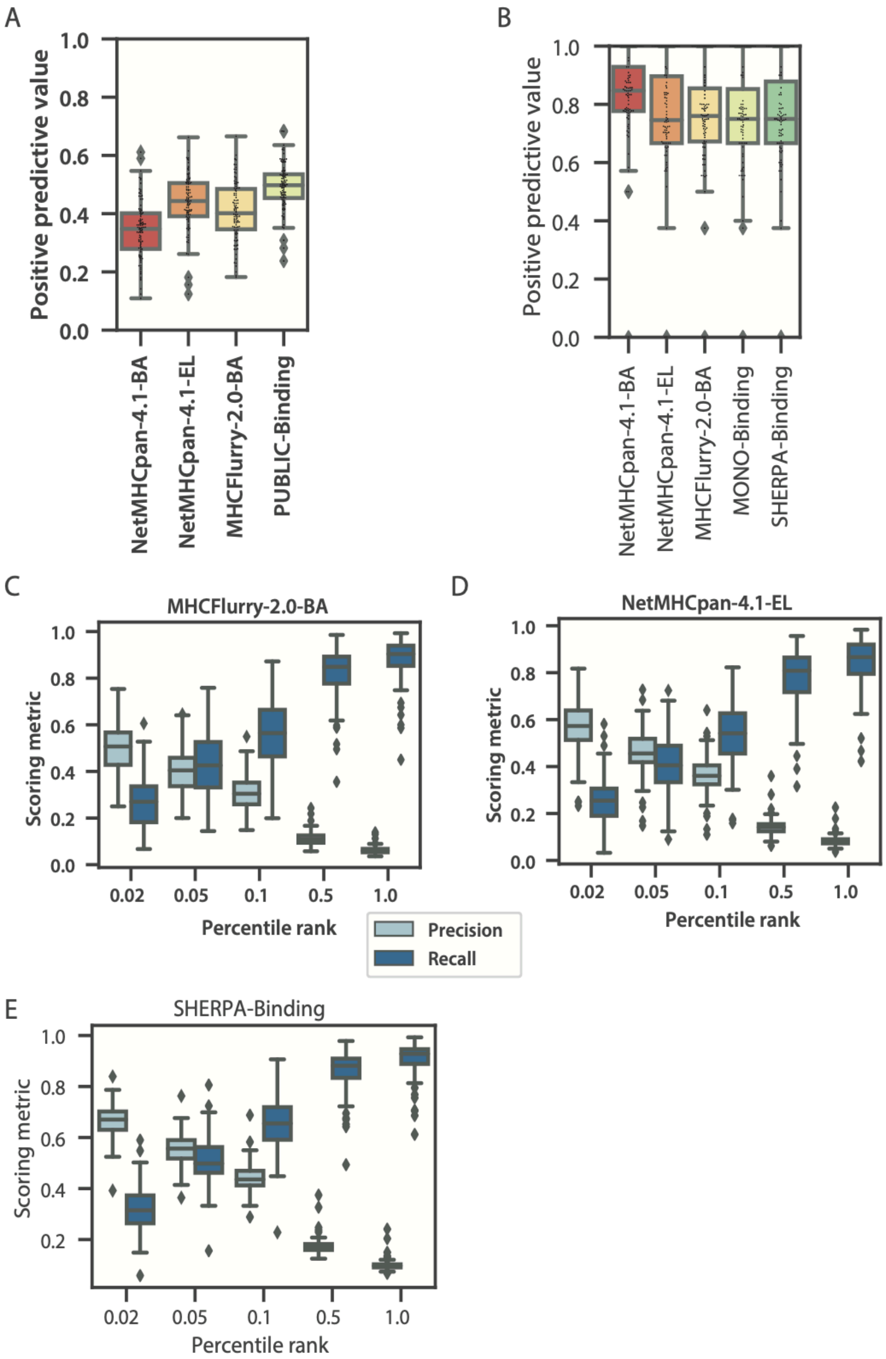


**Supplementary Figure 8: Overview of composite modeling approach and model performance.** (**A**) Boxplots denoting the performance (positive predictive value) of NetMHCpan-4.1-BA, NetMHCpan-4.1-EL, MHCFlurry-2.0-BA and Public-Binding on mono-allelic immunopeptidomics data. (**B**) Boxplots denoting the performance (positive predictive value) of NetMHCpan-4.1-BA, NetMHCpan-4.1-EL, MHCFlurry-2.0-BA, MONO-Binding and SHERPA-Binding on IEDB binding array data. **(C-E)** Boxplots showing the distribution of precision and recall values across alleles in the mono-allelic immunopeptidomics data for (**C**) MHCFlurry-2.0-BA, (**D**) NetMHCpan-4.1-EL and (**E**) SHERPA-Binding across several percentile rank thresholds.

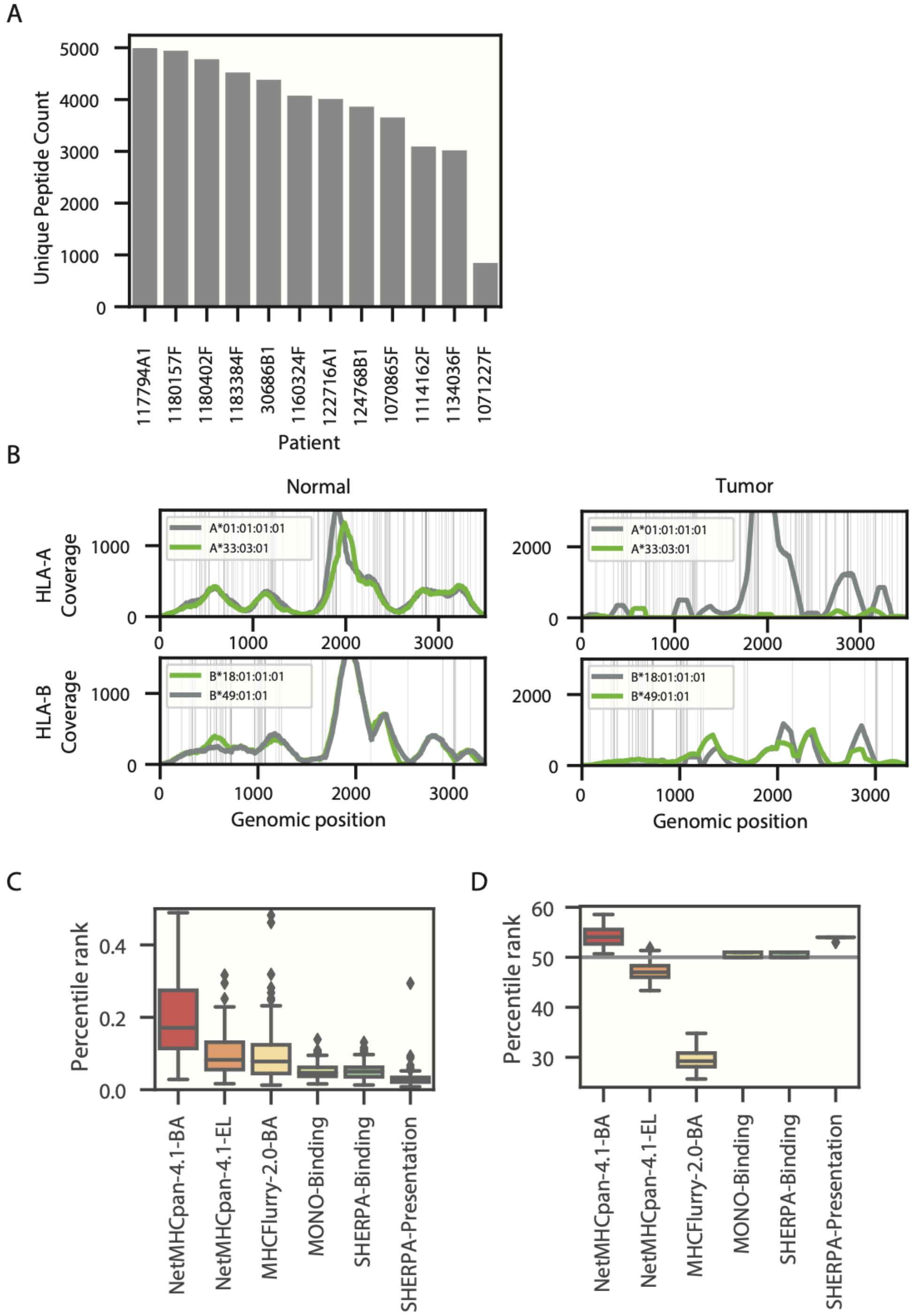


**Supplementary Figure 9:** Overview of tumor immunopeptidomics data and model biases. **(A)** A bar plot denoting the yields of unique peptides from the immunopeptidomics experiments for the 12 tumor samples. **(B)** Line plots showing the sequencing depth of HLA-A and -B alleles in the normal and tumor samples of the patient with HLA LOH and poor performance across all prediction models. (**C-D**) Box plots showing the distribution of percentile ranks for (**C**) positive and (**D**) negative peptides from the mono-allelic dataset.
